# Crosslinking-mediated Interactome Analysis Identified PHD2-HIF1α Interaction Hotspots and the Role of PHD2 in Regulating Protein Neddylation

**DOI:** 10.1101/2024.12.16.628769

**Authors:** Haiping Ouyang, Cindy Y. How, Xiaorong Wang, Clinton Yu, Ang Luo, Lan Huang, Yue Chen

**Author notes:** Correspondence: Dr. Yue Chen or Dr. Lan Huang. indicates equal contributions (Haiping Ouyang, Cindy Y. How).

## Abstract

Prolyl Hydroxylase Domain protein 2 (PHD2) targets Hypoxia Inducible Factor alpha subunits (HIFα) for oxygen-dependent proline hydroxylation that leads to subsequent ubiquitination and degradation of HIFα. In addition to HIF proteins, growing evidence suggested that PHD2 may exert its multifaceted function through hydroxylase-dependent or independent activities. Given the critical role of PHD2 in diverse biological processes, it is important to comprehensively identify potential PHD2 interacting proteins. In this study, we engineered HeLa cells that stably express HTBH-tagged PHD2 to facilitate the identification of PHD2 interactome. Using DSSO-based cross-linking mass spectrometry (XL-MS) technology and LC-MS^n^ analysis, we mapped PHD2-HIF1α interaction hotspots and identified over 300 PHD2 interacting proteins. Furthermore, we validated the COP9 Signalosome (CSN) complex, a major deneddylase complex, as a novel PHD2 interactor. DMOG treatment promoted interaction between PHD2 and CSN complex and enhanced the deneddylase activity of the CSN complex, resulting in increased level of free Cullin and reduced target protein ubiquitination. This mechanism may serve as a negative feedback regulation of the HIF transcription pathway.

## INTRODUCTION

Prolyl Hydroxylase Domain (PHD) proteins are a family of dioxygenases that are highly conserved in metazoan from C. elegans to human (1, 2). They have been extensively characterized for their roles in catalyzing oxygen-dependent hydroxylation on proline residues of hypoxia-inducible factor alpha subunits including HIF1α, HIF2α, HIF3α, which promotes the interaction of HIFα proteins with the von Hippel-Lindau (VHL) subunit of the CUL2 E3 ligase complex and lead to their subsequent polyubiquitination as well as rapid proteasome degradation (3–8). Hypoxia condition significantly reduces the activities of PHD proteins and prevents the proline hydroxylation as well as the rapid degradation of HIFα proteins, which allows the transcriptional activation of hypoxia response genes (9–12). PHD proteins are composed of PHD1 (EGLN2), PHD2 (EGLN1) and PHD3 (EGLN3) as well as a membrane localized P4HTM. Though each PHD protein is capable of catalyzing HIFα proline hydroxylation, PHD2/EGLN1 protein has been known to play a major role in catalyzing HIFα hydroxylation in cells and tissues in response to the changes of physiological concentration of oxygen (13, 14). Genetic loss of PHD2 in mouse was embryonically lethal (15). Interestingly, PHD2 protein is also the transcriptional target of HIF1α, which serves as a negative feedback mechanism to prevent the persistent activation of hypoxia response pathways (16–18).

Among soluble PHD family proteins, PHD2 harbors an additional N-terminal zinc finger zf-MYND domain that facilitates its interaction with other proteins (19, 20). In addition to targeting HIFα proteins for proline hydroxylation, PHD2 has been known to interact with other proteins in hydroxylase-dependent or independent manner. Recent studies showed PHD2 regulates AKT hydroxylation to inhibit its kinase activity, interacts with BRD4 to mediate its interaction with CDK9 in a hydroxylation dependent manner and targets SFMBT1 for pVHL-dependent degradation (21–23). On the other hand, in breast cancer, PHD2 directly interacts with EGFR and stabilizes EGFR levels (24). PHD2 can also form complex with ING4 to inhibit HIF1α transcriptional activities and suppress tumor progression (25). Binding of CIN85 with PHD2 inhibits its activity and promotes cancer progression (26). These studies have shown that PHD2 interacts with a diverse range of proteins, and a comprehensive analysis of its interactome will be essential for identifying novel regulators and substrates, thereby advancing our understanding of its biological functions and significance.Mass spectrometry-based interactome profiling is an effective strategy to identify novel protein-protein interactions. Previous studies have shown that cellular treatment with DMOG, a cell permeable chemical that mimics the PHD2 enzyme cofactor, alpha-ketoglutarate, can further stabilize the PHD2 enzyme-substrate interaction and facilitate the identification of binding targets, a strategy known as the substrate trapping (27). We have previously performed interactome analysis of PHD2 interacting proteins in Hela cells via co-immunoprecipitation and applied label-free quantitative analysis with DMOG-based substrate trapping (28). Our study identified CUL3, a member of the Cullin E3 ligase family, as a novel interactor of PHD2. Functional analysis demonstrated that CUL3-KEAP1 E3 ligase complex regulates PHD2 polyubiquitination and proteasome-mediated protein degradation. Loss of the complex promoted the stabilization of PHD2 under hypoxia and reduced HIF1a abundance (28).

Despite these advances, interactome analysis through traditional co-immunoprecipitation suffers from the loss of sensitivity due to the low affinity in protein-protein interactions and the lack of direct evidence of interactions. XL-MS has been widely used in interactome analysis to enable the capture of native protein interactions and their identification with contact sites at specific amino acid residues (29–31). Classic non-cleavable crosslinkers such as formaldehyde and DSS are capable of stabilizing protein-protein interactions for analysis. However, the resulting cross-linked peptides yield complex MS/MS spectra, making their analysis and unambiguous identification computationally challenging and time consuming (32).

In this study, we applied MS-labile crosslinker DSSO-based XL-MS technology in combination with DMOG-mediated substrate trapping to systematically characterize PHD2 interactome (33). By co-overexpressing HIF1α, we identified endogenous PHD2 interaction sites with HIF1α in solution and label-free quantitative analysis identified 319 binding proteins of PHD2. In addition to canonical PHD2 substrates HIF1α and HIF2α, we identified multiple members of the COP9 Signalosome (CSN) complex, the major deneddylase complex in cells, as novel PHD2 interacting proteins. Functional analysis revealed the DMOG-dependent interaction of PHD2 with the CSN complex promotes the CSN complex activity, reduces the amount of neddylated CUL3 and the polyubiquitination of CUL3 targets. As PHD2 is also a polyubiquitination target of CUL3, such regulation formed negative feedback to control the activation of the HIF transcription-response pathway in cells.

## RESULTS

### Application of MS-cleavable XL-MS to identify PHD2-HIF1**α** interaction sites

MS-cleavable XL-MS has been widely applied in interactome studies to effectively induce stable protein-protein interactions in co-immunoprecipitation and enable confident identification of crosslinking sites(31). To integrate MS-cleavable crosslinking into the workflow of PHD2 interactome analysis, we generated a stable HeLa cell line expressing N-terminal HTBH-tagged PHD2 or empty HTBH tag control through lentiviral transfection (**Figure 1A**). HTBH tag is a multifunctional protein tag optimized for protein complex purification that contains a biotinylation signal (34–36) and therefore, HTBH-tagged PHD2 would be biotinylated in vivo (**Figure 1B**). The Biotin tag allows highly efficient purification of the bait protein with streptavidin-conjugated beads under both native and denatured conditions. We applied this system to identify PHD2-HIF1α interaction sites and map PHD2 interactome. As a central regulator of cellular hypoxia response network, PHD2-dependent interaction of HIF1α has been extensively investigated. However, previous studies mainly focused on crystal structure analysis using recombinant protein domains and synthetic peptides (8, 37, 38). Endogenous interaction sites between full-length PHD2-HIF1α in solution have not been characterized. To this end, we transfected Flag-tagged HIF1α into HeLa cells stably expressing HTBH-tagged PHD2 (**Figure 1C**). After overnight protein expression, cells were treated with 2 mM DMOG for four hours to stabilize PHD2-HIF1α interaction. All cells were lysed with NP-40 lysis buffer by repeated passages through 22G syringe needle. Upon clearing insoluble components, lysates were incubated with streptavidin agarose beads for 2∼3 hours at 4 °C. After washing the beads with lysis buffer and PBS, interacting proteins were crosslinked with 1mM DSSO for one hour followed by quenching with ammonium bicarbonate. Finally, interacting proteins were reduced and alkylated followed by tryptic digestion on beads. Peptides were extracted and desalted for LC MS^n^ analysis to identify DSSO crosslinked peptides (**Figure 1C**).

**Figure 1.**
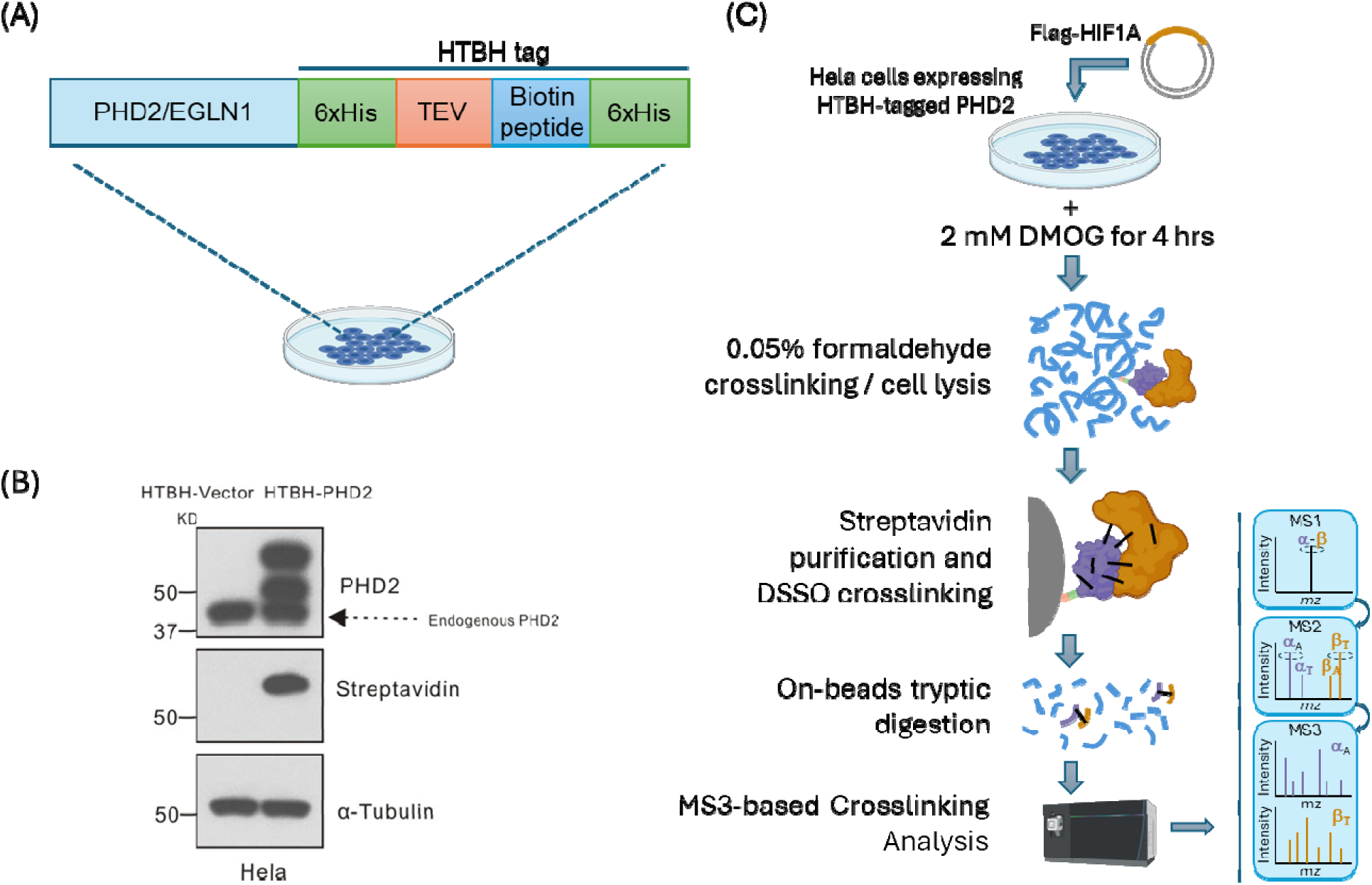
Analytical strategy for crosslinking-mediated identification of PHD2-HIF1α interaction sites. (A) A schematic representation of HTBH-tagged PHD2 for stable expression in HeLa cells. (B) Western blotting validation of HeLa cell lines with or without stable expression of HTBH-tagged PHD2 using PHD2 antibody and streptavidin (SA)-HRP. (C) Analytical workflow to identify in vivo HIF1α-PHD2 crosslinking sites with DSSO crosslinking and DMOG trapping strategies. Crosslinked peptides were identified through MS3 analysis triggered by signature fragmentations in MS2 spectra.

From this analysis, we confidently identified six PHD2-HIF1α cross-linked sites including PHD2/EGLN1 (K146) - HIF1α (K709), PHD2/EGLN1 (K244) - HIF1α (K389), PHD2/EGLN1 (K249) - HIF1α (K389), PHD2/EGLN1 (K291) - HIF1α (K709), PHD2/EGLN1 (K359) - HIF1α (K538), PHD2/EGLN1 (K423) - HIF1α (K709) (**Figure 2**) (**Figure S1, Table S1**). With the full-length PHD2 and HIF1α structures unavailable, this data provides the first glimpse into the mode of endogenous PHD2-HIF1α interaction in solution. From this data, we observed that HIF1α K709 is an interaction hotspot that lies within the close vicinity of K146, K291 and K423 of the PHD2. Available crystal structure of PHD2 only included K291 and K423 but not K146 (7, 8). Using AlphaFold-predicted full-length PHD2, we observed that all three lysine positions lied within the flexible regions of PHD2 with K291 marking the start of the prolyl hydroxylase catalytic domain of PHD2 and K423 lying close to the end of the catalytic domain and protein C-terminus (**Figure S2**). Importantly, HIF1α K709 has been identified as a key acetylation site that is regulated by SIRT2 (39). SIRT2 overexpression promoted HIF1α interaction with PHD2 and subsequent HIF1α degradation. This biochemical analysis corroborated with our finding that K709 of HIF1α is an interaction hub with PHD2 and therefore, acetylation of K709 potentially disrupts the close interaction between HIF1α with multiple regions of PHD2 and reduces PHD2-HIF1α interaction. Our data further identified the three key regions of PHD2 around K146, K291 and K423 that closely interacted with HIF1α K709 (**Figure 2**).

**Figure 2.**
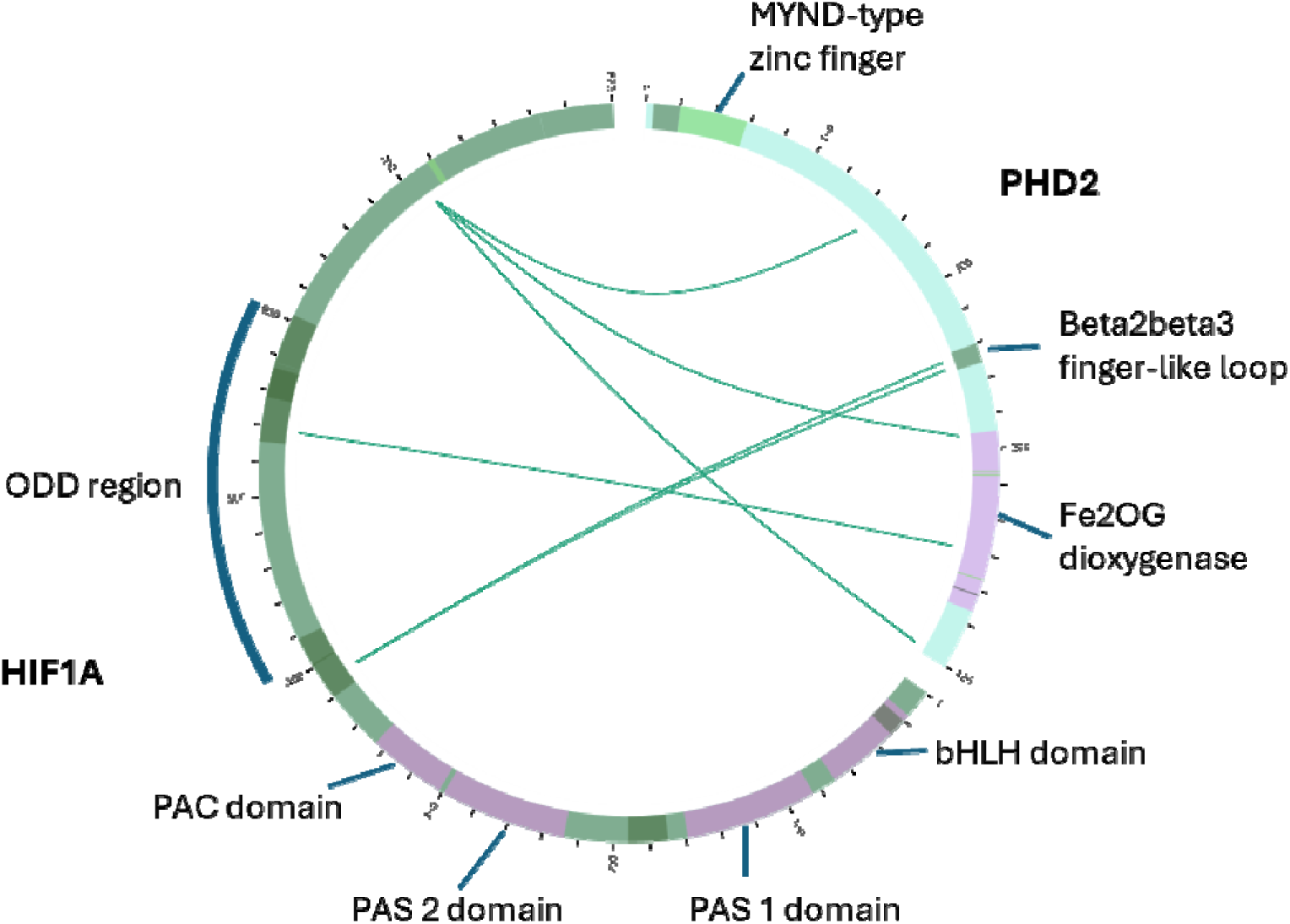
Circular diagram representation of PHD2-HIF1α interaction sites. Full-length UniProt sequences of PHD2 (Q9GZT9) and HIF1α (Q16665) were imported into the xiVew web portal with annotations of domain and regions of interests.

The other two inter-protein cross-linkson HIF1α K389 and K538 were close to the well characterized hydroxyproline modification P402 and P564 on HIF1α that represent N-terminal Oxygen-dependent Degradation Domain (NODD) and C-terminal Oxygen-dependent Degradation Domain (CODD), respectively (**Figure S3**) (40–42). HIF1α K389 was also known to be acetylated and PCAF-mediated acetylation of HIF1α protected it from degradation, potentially through disrupting interactions with PHD2 (43). Interestingly, HIF1α K389 closely interacted with PHD2 K244 and K249, both of which are located on a ten-amino acid β2β3 finger-like loop domain on PHD2 (V241-I251). The β2β3 finger-like loop domain was characterized to be a critical determinant of PHD2 selectivity towards HIF1α hydroxylation on NODD or CODD and the loss of this domain led to preferential PHD2 activity towards CODD (44). Our finding that HIF1α K389 in the NODD closely interacts with β2β3 finger-like loop domain of PHD2 may explain the importance of this domain in maintaining selectivity of PHD2 catalytic activity towards NODD. On the other hand, our data showed that HIF1α K538 in the CODD closely interacted with K359 which locates at the center of the catalytic triad of PHD2 (H313-D315-H374) (8). Such close interaction between HIF1α CODD and PHD2 catalytic center is likely important for efficient CODD hydroxylation and HIF1α degradation.

To integrate the crosslinking data with existing structural information of PHD2 and HIF1α, we applied Alphalink2, an Alphafold-based software application that models protein complex interactions based on known protein structures and identified residue-specific protein crosslinks. After exhaustively analyzing over 200 potential models, we were able to identify top ranked models with scores of above 0.54 and the explanation of one-third of crosslinks (**Figure S4A**). Unfortunately, these top scored models were only able to present the interactions between PHD2 catalytic center and the NODD (P402) of HIF1α (**Figure 3A**). The close distance between HIF1α K389 and the K244, K249 of PHD2 based on the crosslinking data positioned the HIF1α NODD right in the center of PHD2 catalytic center with the P402 site directly facing the PHD2 catalytic triad H313-D315-H374 (**Figure S4B**). From this model, we analyzed the ionic interactions between HIF1α residues and charged surface of PHD2 and we were able to observe a few highly close interactions that likely formed ionic bonds between the side chains and contributed to the stability of the PHD2-HIF1α NODD complex. These interaction hotspots included PHD2 E348 and HIF1α K391 (5.011 Å), PHD2 D246 and HIF1α K389 (10.973 Å), PHD2 K244 and HIF1α E393 (9.419 Å), PHD2 R312 and HIF1α D388 (3.360 Å), PHD2 K402 and HIF1α D395 (6.659 Å), PHD2 R396 and HIF1α D406 (7.624 Å), PHD2 D212 and HIF1α K362 (2.697 Å), PHD2 K204 and HIF1α D368 (2.669 Å) (**Figure 3B-D**). Further biochemical studies are warranted to investigate the functional significance of these interactions in full-length PHD2-HIF1α complex stability in vivo.

**Figure 3.**
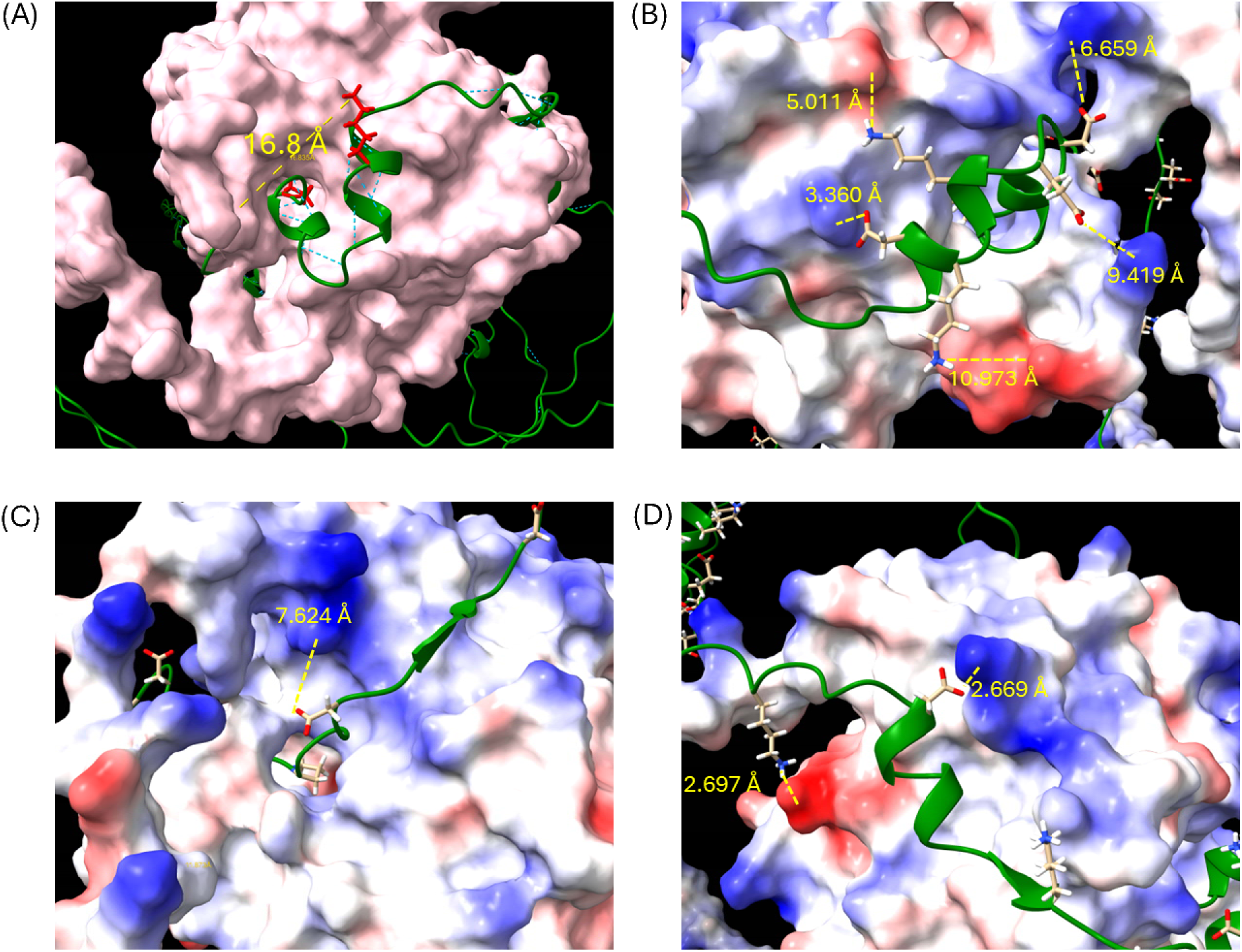
Structural representations of Alphalink2-predicted interaction between HIF1α NODD region with PHD2 catalytic center. (A) Predicted domain interaction between PHD2 catalytic center and HIF1α NODD region (in green) with HIF1α K389 – PHD2 K244 distance highlighted in yellow. Structural representation of potential ionic interactions between PHD2 catalytic center and HIF1α NODD based on Alphalinks2 predictions with (B) PHD2 E348 and HIF1α K391 (5.011 Å), PHD2 D246 and HIF1α K389 (10.973 Å), PHD2 K244 and HIF1α E393 (9.419 Å), PHD2 R312 and HIF1α D388 (3.360 Å), PHD2 K402 and HIF1α D395 (6.659 Å), (C) PHD2 R396 and HIF1α D406 (7.624 Å), (D) PHD2 D212 and HIF1α K362 (2.697 Å), PHD2 K204 and HIF1α D368 (2.669 Å).

### Crosslinking-assisted system-wide profiling of PHD2 interactome

Next, we applied the MS-cleavable crosslinking technology and DMOG-mediated substrate trapping for system-wide profile PHD2 interacting proteins in Hela cell lines with label-free quantitative analysis. To this end, HeLa cells stably expressing control plasmid or HTBH tagged PHD2 were cultured in triplicates. An additional set of HeLa cells expressing HTBH-PHD2 were treated with 2 mM of DMOG for four hours to induce substrate trapping. Cells were lysed and subject to affinity purification, crosslinking and on-beads tryptic digestion HIF1αprior to LC MS^n^ and LC MS/MS analysis respectively (**Figure 4A**).

**Figure 4.**
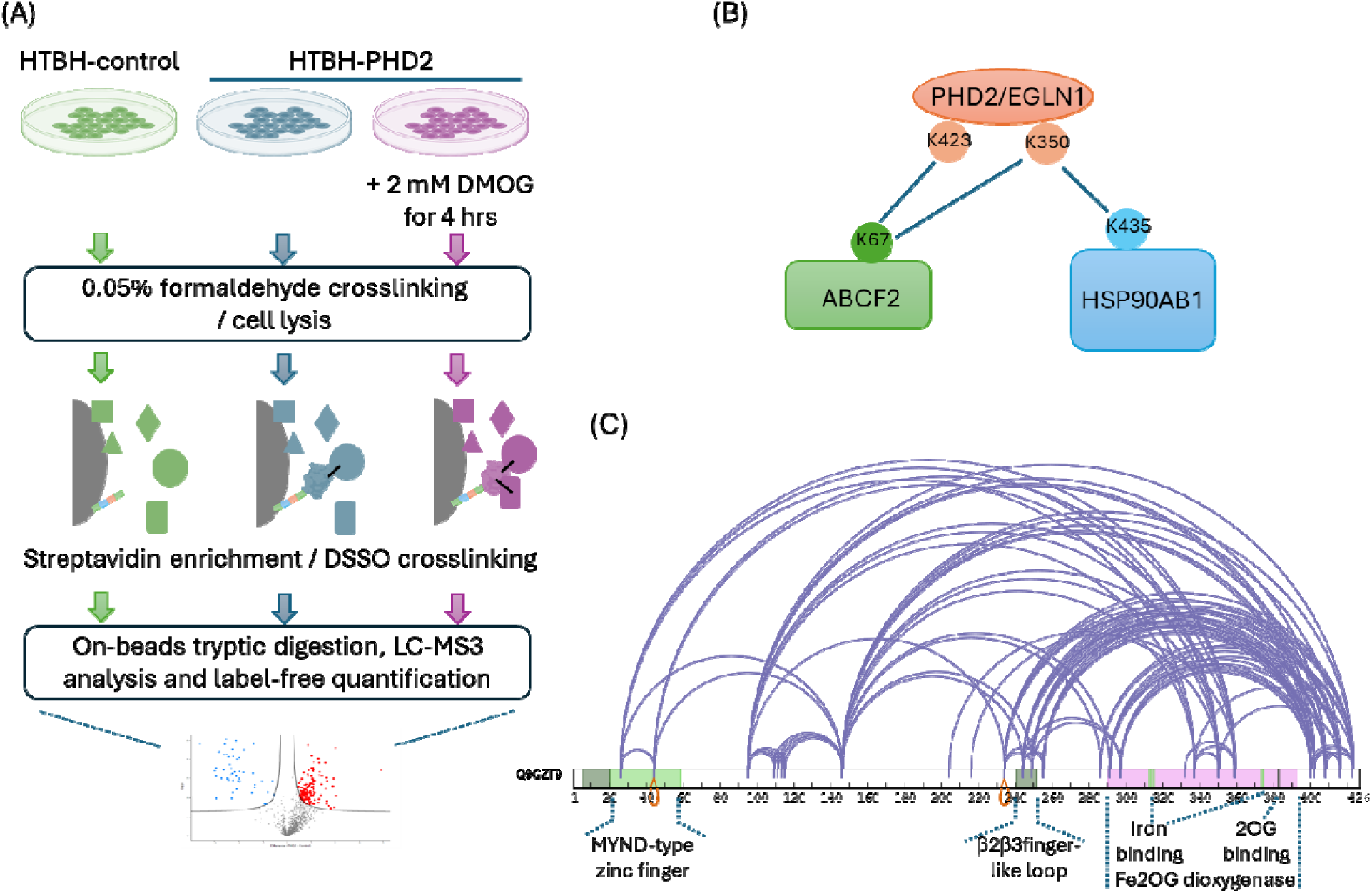
Crosslinking-mediated PHD2 interactome analysis. (A) Analytical workflow to identify PHD2 interactome with HeLa cells expressing HTBH-control and HTBH PHD2 with or without DMOG trapping. (B) Identification of endogenous PHD2 intercrosslinks with ABCF2 and HSP90AB1. (C) Identification of endogenous PHD2 intracrosslinks. Full length sequence of UniProt PHD2 (Q9GZT9) was imported into the xiView web portal with annotations of domain and regions of interests.

From this analysis, we identified 142 intralinks and interlinks of PHD2 (**Table S1**). Due to the high background of system-wide analysis, we only identified three PHD2 interlinks: ABCF2 (K67) ∼ PHD2/EGLN1 (K350), ABCF2 (K67) ∼ PHD2/EGLN1 (K423) and PHD2/EGLN1 (K350) ∼ HSP90AB1 (K435) (**Figure 4B**). ABCF2 belongs to the ATP-Binding Cassette (ABC) transporter superfamily but has no transmembrane domain. It was not previously known to interact with PHD2 and the functional significance of such interaction remains to be investigated. PHD2 has been previously shown to interact with HSP90 complex including HSP90, cochaperone p23 and FKBP38 (19). Our data revealed the close interaction between HSP90AB1 and PHD2 catalytic domain residue K350. Among intralinks, we identified 99 linkages on PHD2 and the most frequently identified linkages included K291∼K423, K249∼K402, K146∼K402 and K146∼K350. Importantly, we observed that multiple adjacent residues on distal domains of PHD2 formed dense clusters of linkages (**Figure 4C**). Previously, PHD2 has been reported to mostly exist in monomer form in solution (7, 45). Therefore, our data presented a high confidence map of full-length PHD2 domain connectivity that may link to regulatory functions.

### Quantitative interactome analysis identified the PHD2-CSN complex interaction

For quantitative analysis of PHD2 interactome, MS data was then processed with Maxquant and Perseus computational platform and Intensity-Based Absolute Quantification (IBAQ) was calculated for label-free quantification of PHD2 interacting proteins (46–48). Without DMOG treatment, we identified 289 proteins that significantly interacted with PHD2 and 158 proteins with DMOG treatment (**Figure 5A and S5, Table S2**). Among them, 128 proteins overlapped between the two studies and showed more stable interaction with PHD2 regardless of DMOG treatment (**Figure 5A**). Gene Ontology enrichment analysis showed that the proteins that stably interacted with PHD2 were significantly enriched in biological processes including protein localization to ER, RNA catabolic process and cytosolic translation (**Figure 5B**).

**Figure 5.**
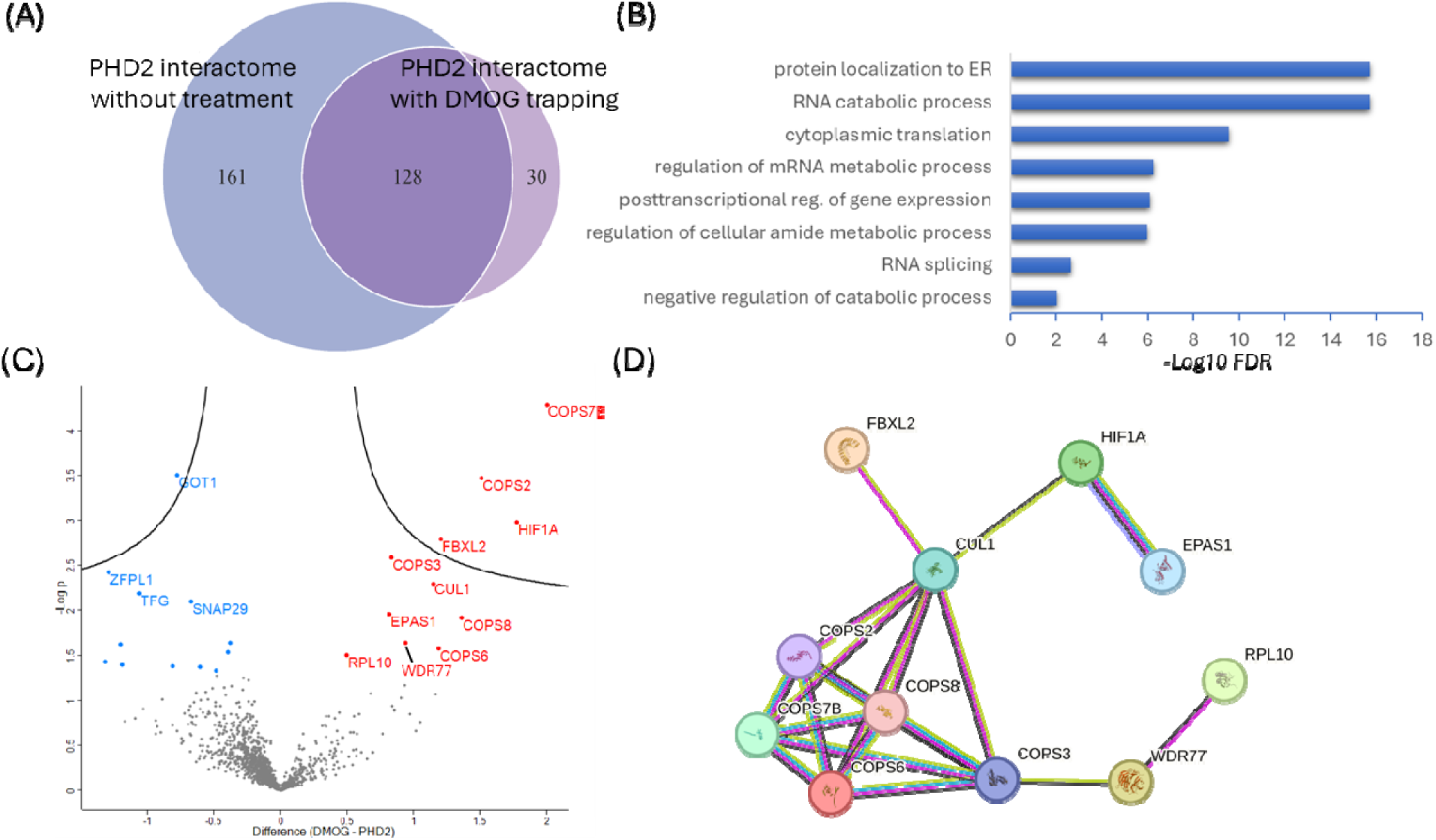
Quantitative profiling PHD2 interactome. (A) Venn diagram representing the overlap of PHD2 interacting proteins identified with and without DMOG treatment with label-free quantitative analysis. (B) Annotation enrichment analysis of Gene Ontology biological processes with FDR cutoff of 0.05. (C) Volcano plot of the label-free quantitative comparison of PHD2 interacting proteins with and without DMOG treatment (FDR<0.05) with selected genes highlighted in red (upregulation with DMOG treatment) and in blue (downregulation with DMOG treatment). (D) The interacting protein network of PHD2 interacting proteins that were upregulated after DMOG treatment. The interactions were extracted from STRING database through web portal.

We further compared the PHD2 interactome with or without DMOG treatment. Among the few proteins having preferential interactions with PHD2 under DMOG treatment were well known PHD2 targets HIF1α and HIF2α (**Figure 5C**). Interaction network analysis showed that these proteins formed a highly connected network that contained a subnetwork composed of five members of the COP9 Signalosome (CSN) Complex (**Figure 5D**). CSN complex is an essential protein complex that catalyzes deneddylation reaction in cells. Neddylation is an important activation mechanism to enhance the assembly and activities of Cullin ubiquitin E3 ligases, the largest family of ubiquitin E3 ligases (49, 50). Therefore, CSN complex plays critical roles in the regulation of ubiquitination pathways by regulating neddylation level on the Cullin ubiquitin E3 ligases (51, 52). Our interactome data suggested that CSN complex may be an interacting partner of PHD2 and their interactions were further stimulated by the DMOG treatment.

To validate this finding, we performed affinity purification and western blotting analysis. Our data clearly showed that PHD2 was sufficient to pulldown multiple members of the CSN complex including CSN2, CSN3, CSN4, CSN7 and CSN9 (**Figure 6A**). While DMOG treatment did not affect the protein abundance of the CSN complex, it strengthened the interaction between PHD2 and the CSN complex. To corroborate with this data, we performed reciprocal affinity purification. 293T cells were transfected with Flag-tagged CSN6 followed by co-immunoprecipitation. Our western blotting clearly confirmed that PHD2 interacted with CSN6 and the interaction became stronger with DMOG treatment (**Figure 6B**). To determine if the interaction between PHD2 and CSN complex is dependent on HIF1α, we performed siRNA knockdown and repeated the co-immunoprecipitation of PHD2 in the stable HeLa cell line expressing HTBH-tagged PHD2. Our data confirmed that PHD2-CSN complex interaction was independent of HIF1α abundance (**Figure S6**). As DMOG treatment has often been applied to mimic the activation of hypoxia response in cells, we tested whether hypoxia treatment can also boost the interaction of PHD2 and CSN complex. The experiment clearly confirmed the PHD2 interaction with CSN complex, but the interaction was not enhanced under hypoxia condition (**Figure S7**). To further validate the findings in other cell lines, we performed transient overexpression of HA-tagged PHD2 in 293T cells. Our data confirmed the PHD2-CSN complex interaction again but the DMOG trapping effect became less apparent when PHD2 protein was overly expressed (**Figure S8**).

**Figure 6.**
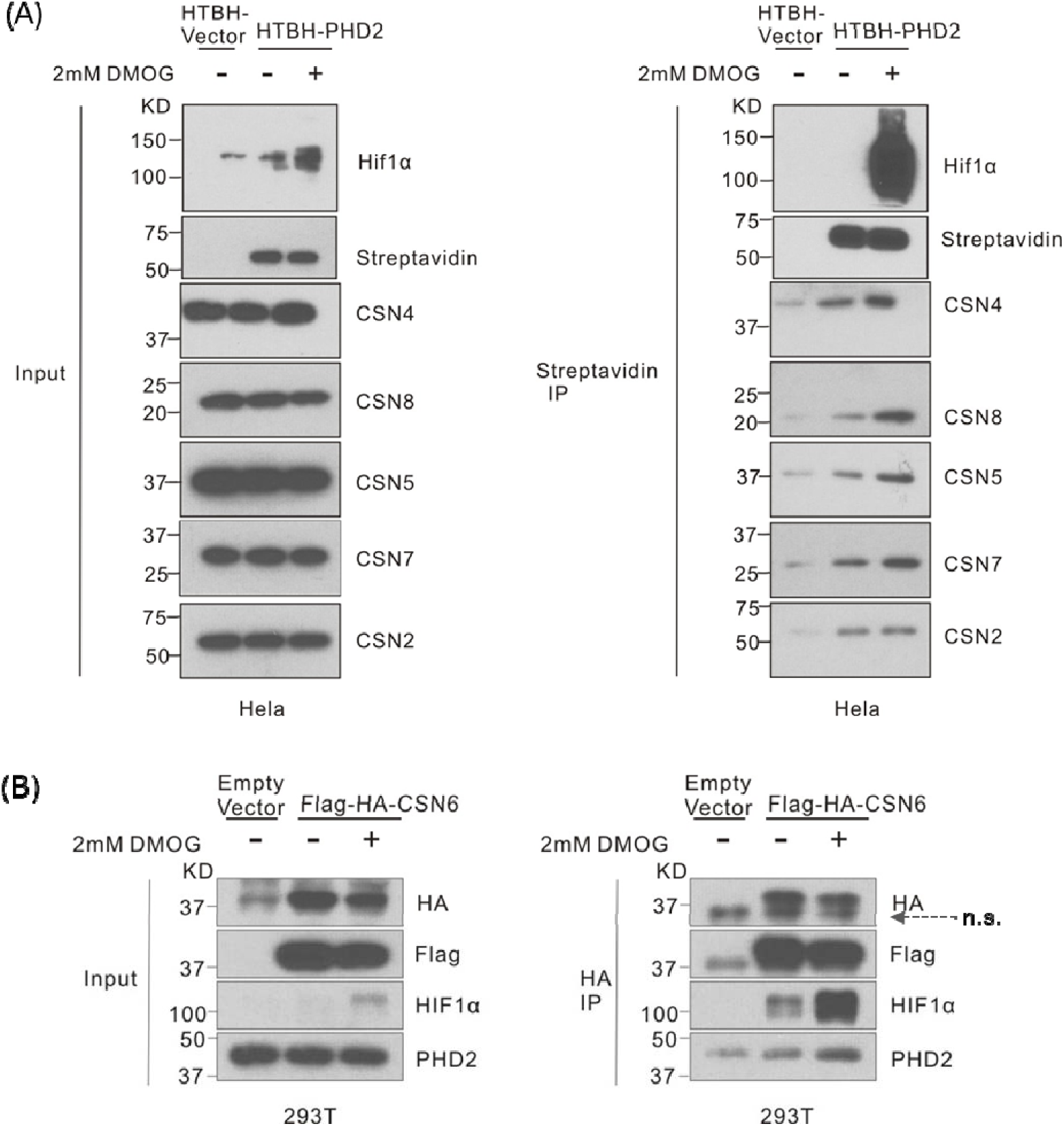
Validation of PHD2 interaction with the CSN complex. (A) Western blotting validation of CSN complex members interaction with PHD2 in stable HeLa cell lines. Hela cells that stably express HTBH-empty vector or HTBH-PHD2 were seeded and treated with DMSO or 2mM DMOG for 4 hours prior to lysis and streptavidin enrichment. The same amount of input and IP samples of each group are loaded for WB analysis and blotted with antibodies as indicated. The expression of HIF1α was examined as the indicator of DMOG treatment. (B) Western blotting validation of PHD2 interaction with CSN6 in 293T cells. 293T cells were transiently transfected with Flag-HA-CSN6 plasmid or empty vector for 18 hours and treated with 2mM DMOG or DMSO for 4hours prior to lysis and immunoprecipitation with anti-HA antibody conjugated beads. The same amount of input and IP samples of each group are loaded for WB analysis and blotted with antibodies as indicated. The expression of HIF1α was examined as the indicator of DMOG treatment. “n.s.” indicates non-specific band.

### Interaction of PHD2 and CSN complex promotes CSN deneddylase activity and reduced the ubiquitination activities of the Cullin E3 ligase families

Next, we investigated the functional significance of PHD2-CSN complex in regulating CSN deneddylase activity. To this end, we performed DMOG treatment to promote PHD2-CSN complex interaction in stable HeLa cells expressing HTBH-PHD2. Neddylation abundance was monitored with western blotting against Nedd8 and several Cullinsincluding CUL1, CUL3 and CUL4A. Our data showed that DMOG treatment reduced neddylation globally and on the selected Cullins (**Figure 7A**). We further confirmed the findings in 293T cells (**Figure 7B**). Several members of the CSN complex have known proline hydroxylation sites including CSN2 (P05814) P180, CSN3 (Q9UNS2) P331, CSN6 (E7EM64) P129, P130, CSN1 (A0A096LP07) P22 and CSN9 (Q8WXC6) P15 (**Figure S9A-F**) (53). It was likely that PHD2 targeted CSN complex for proline hydroxylation, which may regulate the enzymatic activities of the complex. Therefore, it is important to determine whether DMOG regulated neddylation through PHD2 interaction with CSN complex or through prolyl hydroxylase-mediated proline hydroxylation. To this end, we performed siRNA knockdown of PHD2 in 293T cells. Our data clearly showed that the knockdown of PHD2 led to an apparent increase in neddylation levels of CUL1 and CUL3 in vivo (**Figure 7C**). Furthermore, knockdown of PHD2 restored the DMOG-mediated reduction of the neddylation level of CUL3 in 293T cells (**Figure 7D**). As DMOG is a strong inhibitor of all prolyl hydroxylase catalytic activities, if DMOG exerted its effect on cullin neddylation through CSN proline hydroxylation, the knockdown of PHD2 should not affect DMOG-mediated downregulation of neddylation. Therefore, our data strongly suggested that it was PHD2 protein instead of CSN proline hydroxylation playing a critical role for modulating CSN deneddylase activities.

**Figure 7.**
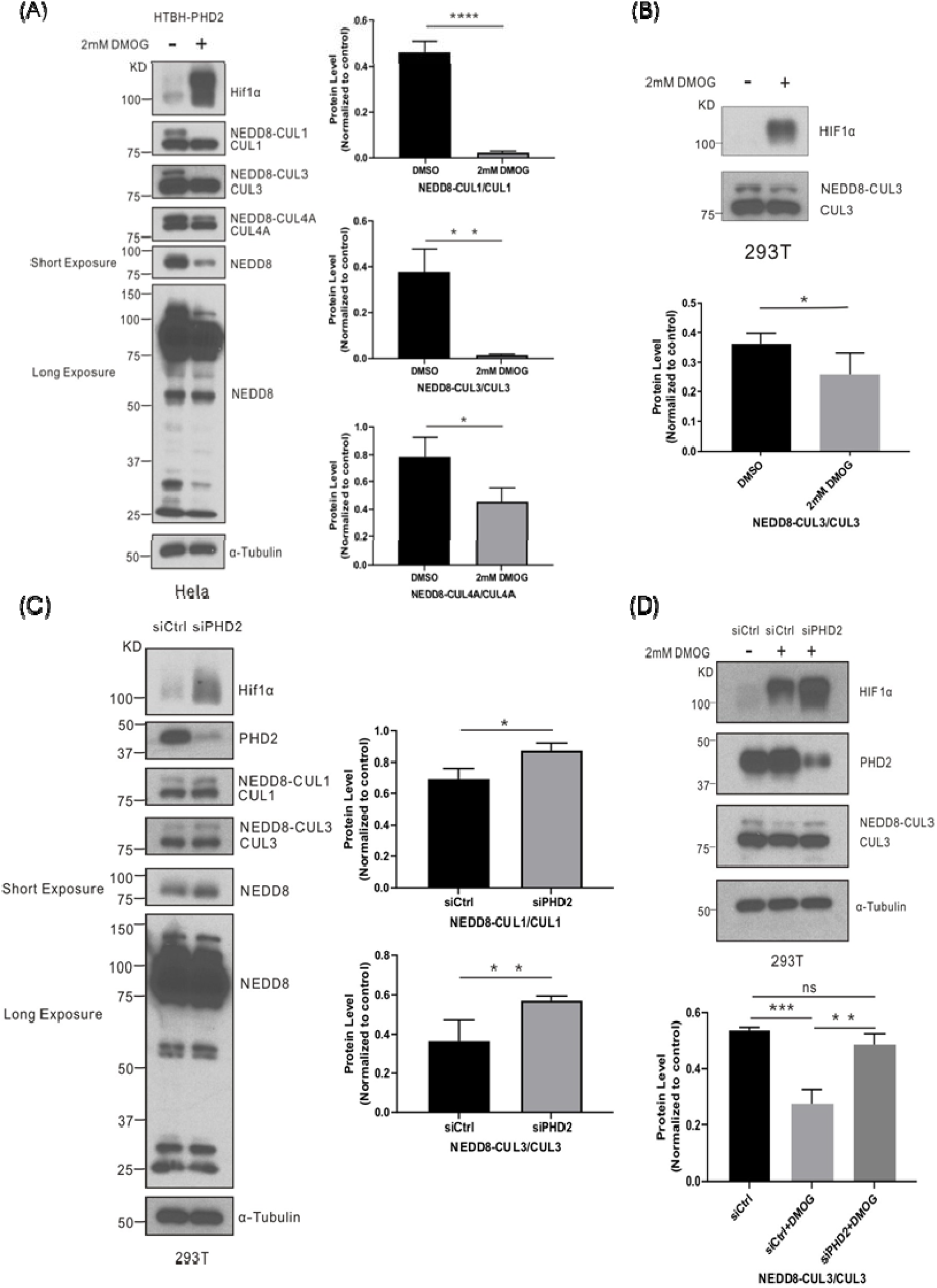
PHD2 interaction with the CSN complex regulates protein neddylation. (A) Western blotting analysis showing the DMOG treatment in HeLa cells significantly downregulated protein neddylation. Hela cells stably expressing HTBH-control were seeded and treated with DMSO or 2mM DMOG for 24 hours under normoxia condition, then cells were harvested and lysed for WB analysis and blotted with antibodies as indicated. The expression of HIF1α was examined as the indicator of DMOG treatment. (B) Western blotting analysis showing that DMOG treatment in 293T cells downregulated Cullin3 neddylation. 293T cells were seeded and treated with DMSO or 2mM DMOG for the next 24 hours and cultured under normoxia condition, harvested for WB analysis. The neddylated CUL3 level was normalized to the total CUL3 expression level. (C) Western blotting analysis showing that siRNA knockdown of PHD2 upregulated Cullin neddylation. 293T cells were seeded and treated with 60nM siRNA targeting PHD2 for 48 hours under normoxia condition, then cells were harvested and lysis for WB analysis and blotted with antibodies as indicated. The expression of HIF1α was examined as the indicator of PHD2 knockdown. (D) Western blotting analysis showing that siRNA knockdown of PHD2 rescued DMOG-mediated downregulation of Cullin3 neddylation. 293T cells were seeded and knockdown of PHD2 by applying 60nM siRNA targeting PHD2 for 18 hours followed by DMSO or 2mM DMOG treatment for the next 24 hours. Then cells are harvested and lysed for WB analysis and blotted with antibodies as indicated. The expression of HIF1α was examined as the indicator of DMOG treatment. n≥3, *P<0.05, ** P<0.01, ***P<0.001.

Stimulation of CSN deneddylase activity would lead to the reduction of Cullin ubiquitin E3 ligase activities. To demonstrate this, we would like to determine if DMOG-mediated changes in Cullin neddylation level may affect the ubiquitination abundance of known Cullin E3 ligase targets. We selected PHD2 and NRF2 as examples, both of which have been previously characterized as the targets of Cullin3 E3 ligase complex (28, 54–56). Western blotting analysis showed that DMOG treatment indeed reduced ubiquitination levels of PHD2 and NRF2 in 293T cells or HeLa cells with or without expressing HTBH-tagged PHD2 (**Figure S10A-B**). This data confirmed that DMOG-mediated decrease in Cullin neddylation led to reduced Cullin E3 ligase activities and reduced polyubiquitination of Cullin substrates.

## DISCUSSION

Prolyl hydroxylases are essential metabolic sensors governing diverse cellular pathways in response to nutrient and oxygen levels in cellular microenvironment. These proteins exert their cellular functions through hydroxylase-dependent or independent roles. Confident and sensitive identification of prolyl hydroxylase interactome lays the foundation of discovering novel PHD-dependent regulation of cellular pathways and physiology. In this study, we reported the chemical crosslinking-mediated quantitative analysis of PHD2 interactome. Integrated with DMOG-mediated substrate trapping and label-free quantitative analysis, our study identified 289 PHD2 interacting proteins without DMOG treatment and 158 PHD2 interacting proteins with DMOG treatment, including many known and novel interacting targets.

XL-MS is an efficient strategy to determine spatial distance of proteins in solution. It is ideally applied to map endogenous interactions in complex biological solution that are typically not available for measurement with other analytical methods such as X-ray, Cryo-EM or NMR. In this study, we applied MS-cleavable XL-MS technology and generated the first map of interprotein linkages between endogenous PHD2 and HIF1α in solution, which not only validated previous biochemical studies but also revealed new insights into the mechanisms of substrate domain recognition. Extensive intra protein linkages of PHD2 identified in this study presented a confident map of interconnected domains and such connections are potentially important in PHD2 enzymatic activity and substrate interaction. It is important to note that these domain interactions were likely connected through flexible linker regions and therefore not easily captured by conventional structural analysis or modeling simulation. Therefore, XL-MS analysis offers additional mechanistic insights on PHD2 domain interactions and substrate binding. Overall, our chemical crosslinking-based interactome analysis pipeline offers new approaches to confidently identify protein interactome that can be generally applied to study other prolyl hydroxylases and identify new regulatory pathways.

In this study, we identified and validated endogenous interactions between PHD2 and COP9 Signalosome (CSN) complex. Importantly, we determined that PHD2-CSN complex interaction enhanced the deneddylase activity of CSN complex to modulate cullin E3 ligases and protein ubiquitination. As the largest family of ubiquitin E3 ligases, such regulation likely plays a critical role for the changes of ubiquitination level and degradation profiles of diverse protein targets upon the activation of the HIF pathway. Interestingly, PHD2 itself is subject to CUL3 mediated ubiquitination and protein degradation. Upon DMOG treatment, HIF pathway is activated. To prevent persistent activation of HIF pathway, HIF1α promotes the transcriptional activation of PHD2 which serves as a negative feedback mechanism to reduce HIF1α abundance and HIF pathway activities. As a synergistic effort, our study showed that DMOG treatment promoted interaction of PHD2 with CSN complex, reduced the neddylation level of CUL3 E3 ligase and reduced ubiquitination of PHD2. Such mechanism cooperated with the HIF1α-mediated increase in PHD2 transcription to synergistically increase PHD2 abundance for negative feedback regulation of HIF pathway activities in cells.

## MATERIALS AND METHOD

### Chemicals and Reagents

DMOG (A4506) was from ApexBio (Houston, TX). Formaldehyde solution (252549), puromycin (540222), NP-40 (I8896), MgCl2 (7791-18-6), Roche cOmplete protease inhibitor cocktail tablet (11697498001), PhosSTOP phosphatase inhibitor (4906845001), Fetal Bovine Serum (FBS) (F0926), polyethyleneimine (PEI) (MilliporeSigma, Burlington, MA), Luminata Crescendo Western HRP Substrate (WBLUR0500) and anti-HA agarose beads (A2095) were from MilliporeSigma (Burlington, MA). Glycine (15527013), glycerol (G33-1), DTT (PI20290), TCEP (PG82080), streptavidin resins (PI20347), LCMS-grade acetonitrile (51101) and water (51140), Dulbecco’s Modified Eagle’s medium (DMEM) (Gibco, 11965092), reduced serum medium (Opti-MEM, 31985070), Bradford assay kit (23200) and DSSO-disuccinimidyl sulfoxide (PIA33545) were from ThermoFisher Scientific (Waltham, MA). Polybrene reagent (sc-134220) was from Santa Cruz BioTech (Dallas, TX). ATP (BML-EW9805-0100) was from Enzo Life Sciences (Farmingdale, NY). Iodoacetamide (02327) was from Chem-Impex (Wood Dale, IL). Difco skim milk was from VWR (90002-594). 100X Penicillin-streptomycin (25–512) was from Genesee Scientific (Morrisville, NC). Sequencing-grade trypsin (V5113) was from Promega (Madison, WI). Anti-HA (660002) antibody and Streptavidin HRP (405210) were from Biolegend (San Diego, CA). Anti-HIF1α (SAB2702132), anti-α-Tubulin (T6199), anti-HA (SAB4300603) and anti-Flag (F1804) antibodies were from MilliporeSigma (Burlington, MA). Anti-CSN2 (10969-2-AP), anti-CSN4 (10464-1-AP), anti-CSN5 (27511-1-AP) and anti-CSN8 (10089-2-AP) antibodies were from proteintech (Rosemont, IL). Anti-CSN7(A300-240A) antibody was from Bethyl (Montgomery, TX). Anti-NEDD8(2754T), anti-CUL1(4995S), anti-CUL3 (2759S), anti-CUL4A (CSIG-2699T), anti-PHD2 (4835), anti-NRF2 (12721T) antibodies and HRP-linked secondary antibody (7076 and 7074) were from Cell Signaling Technology (Danvers, MA). Mycoplasma PCR detection kit (G238) was from Applied Biological Materials (Richmond, British Columbia, Canada)

### Generation of stable HeLa cell lines

The pQCXIP-HTBH-PHD2 plasmid was generated by cloning the CDS sequence of PHD2 into the pQCXIP-HTBH empty vector (34). To establish stable cell lines, 293FT cells were seeded in DMEM medium containing 10% FBS (without antibiotics) to reach around 70% confluence on the next day. Then, HTBH-PHD2 or HTBH-empty vector plasmid were co-transfected into 293FT cells together with pCMV-DR8.2 and pCMV-VSV-G at the ratio of 1:1:0.2 by using polyethylenimine (PEI). Change medium within 18 hours with fresh growth medium. 48 hours after transfection, cell medium containing the lentivirus was filtered with a 0.45 μm filter, and then used to infect Hela cells (ATCC #CCL-2) with the help of polybrene (8 μg/ml). 24 hours after lentivirus infection, the cells were transferred into medium containing 2 μg/ml puromycin for one week. After selection, the positive cells were maintained in medium containing 1μg/ml puromycin and expanded for reservation and analysis.

### Cell culture

HEK293T cells (ATCC #CRL-3216) were cultured in Dulbecco’s Modified Eagle’s medium (DMEM) supplemented with 10% FBS and 1% penicillin-streptomycin. Hela cells that stably express HTBH-empty vector or HTBH-PHD2 were cultured in DMEM supplied with 1ug/ml puromycin, 10% FBS and 1% penicillin-streptomycin. The cells were maintained at 37 °C in incubators supplied with 5% CO_2_. For hypoxia treatment, cells were cultured in a CO_2_ incubator supplied with 1%O_2_ / 94% N_2_ / 5% CO_2_. Cell lines were regularly monitored for mycoplasma to ensure no contamination.

### Transient transfection for overexpression

The pc-DNA plasmid vectors containing HA-tagged PHD2 was a kind gift from Do-Hyung Kim (University of Minnesota). HA-Flag-tagged CSN6 plasmid was a kind gift from Wade Harper (Addgene #22542). Plasmid transfection in HEK293T cells was performed using 1 mg/ml polyethyleneimine. Transfection was conducted with a 1:3 ratio (μg DNA: μg PEI) in reduced serum medium (Opti-MEM) one day after cell seeding.

### Gene knockdown with siRNA

Transfection with HIF1A siRNA (hs.Ri.HIF1A.13.3, dsiRNA targeting HIF1A, NM_001243084, NM_181054, NM_001530, CDS-Exon 9) (Integrated DNA Technologies, Coralville, Iowa), PHD2 siRNA (5′-GACGAAAGCCAUGGUUGCUUG-3′ (22)) or control siRNA (SIC001, MISSION® siRNA Universal Negative Control) (MilliporeSigma, Burlington, MA) was performed with DharmaFECT 1 Transfection Reagent according to the manufacturer’s instructions (T-2001-02) (Thermofisher Scientific, Waltham, MA).

### Enrichment and co-immunoprecipitation analysis with chemical crosslinking and western blotting

Hela cells that stably expressed HTBH-empty vector or HTBH-PHD2 with or without transfection of Flag-HIF1α were seeded and treated with DMSO or 2 mM DMOG for 4 hours, then crosslinked with 0.1%formaldehyde solution for 10 minutes at 37°C followed by quenching with 125mM glycine for 5 minutes at 37°C. Then cells were washed with pre-cold PBS and lysed with pre-cold lysis buffer (0.5% NP-40, 10% glycerol, 1 mM ATP, 1 mM DTT, 5 mM MgCl_2_, 2 mM DMOG, supplied with 1X cOmplete protease inhibitor and 1X phosphatase inhibitor). Cell lysate was passed through a 22G needle 20 times to completely lysis the cells then centrifuge at 18000 g for 15 minutes at 4 °C to remove cells debris. Protein concentration in the clear cell lysate was measured using Bradford assay. About 5% (250 μg) of protein was used as input for western blotting and 5 mg protein was incubated with streptavidin resins at 4°C for 2 hours with rotation. Subsequently, the streptavidin resin conjugated protein complex was spin down at 800xg at 4°C for 1 min followed by washing with pre-cold lysis buffer once and pre-cold PBS once to remove non-specific binding.

For reciprocal Co-IP, 293 T cells were grown to 60% confluence and transiently transfected with HA-Flag-tagged empty vector or HA-Flag-tagged CSN6 expression plasmids. Sixteen hours after transfection, cells were treated with DMSO or 2mM DMOG for 4 hours. Then formaldehyde crosslinking and cell lysis were done as mentioned above. To conduct co-immunoprecipitation, cell lysates were incubated with anti-HA agarose beads at 4 °C overnight followed by washing with pre-cold lysis buffer once and pre-cold PBS once to remove non-specific binding prior to elution.

For western blotting detection, elution was performed with SDS sample buffer (375 mM Tris-HCl, pH 6.8, 9% SDS, 50% glycerol, 6% β-mercaptoethanol, 0.03% bromophenol blue) and boiled at 99°C for 10 minutes. Equal amounts of input and immunoprecipitated protein were separated by SDS-PAGE, transferred to PVDF membranes (1620177) (Bio-Rad, Hercules, CA) and incubated with antibodies as indicated. Detailed steps of western blotting were described below.

### Western blotting

For western blot assays in cell lysates, after the treatment as indicated, cells were harvested by washing with pre-cold PBS and lysed in pre-cold lysis buffer (150 mM NaCl, 0.5% NP-40, 50 mM Tris-HCl, 10% glycerol, 1% SDS, pH 7.5, supplemented with 1x cOmplete protease inhibitor cocktail) on ice for 15 min. After removal of cell debris by centrifugation at 21,000xg at 4°C for 10 min, the soluble fractions were collected and boiled at 99°C in SDS sample buffer for 10 min. The extracted proteins were resolved in homemade SDS-PAGE gel and transferred onto PVDF membrane. Blocking was done with 5% skim milk in TBST (TBS+0.1% Tween-20) for 1 hour. After blocking, the membrane was incubated with primary antibodies as indicated overnight at 4°C and washed with TBST for 3 times before 1 hour incubation with HRP-linked secondary antibody at room temperature. The signal was developed with Luminata Crescendo Western HRP Substrate. ImageJ software was used to quantify the protein bands. For western blot quantification, data are represented as the mean+-S.E.M of three independent experiments. Statistical analyses were performed in GraphPad Prism with the Student’s t-test.

### DSSO crosslinking and proteomic sample preparation

PHD2 and its interacting proteins were precipitated on streptavidin beads from stable Hela cells expressing HTBH-PHD2 or control with or without DMOG treatment or the transfection of Flag-HIF1α as described above. After washing, the beads were resuspended in 1mM DSSO MS-cleavable crosslinking reagent in 1X PBS and incubated at room temperature for one hour. The beads were washed twice with 25 mM NH_4_HCO_3_ buffer with rotation for one minute each time to stop crosslinking. The beads were then resuspended with 2 mM TCEP and 10 mM IAA for 30 minutes in dark at room temperature with rotation. For enzymatic digestion, each batch of beads was washed once with NH_4_HCO_3_ buffer and then resuspended in the NH_4_HCO_3_ buffer with 1.5 M urea. Two micrograms of trypsin were added to each batch of beads for digestion overnight at 37°C and pH 7. The tryptic peptides in solution were then extracted, desalted with in-house packed C18 Stage Tip, and eluted with 50% acetonitrile. Elutes were dried with SpeedVac Vacuum Concentrator (ThermoFisher Scientific, Waltham, MA) prior to the XL-MS analysis.

### LC-MS^n^ analysis and identification of DSSO-crosslinked peptides

LC-MS^n^ analysis of cross-linked peptides was performed using an UltiMate 3000 UPLC (Thermo Fisher Scientific) liquid chromatograph coupled on-line to an Orbitrap Fusion Lumos mass spectrometer (Thermo Fisher Scientific). Peptides were separated by reverse-phase on a 50cm x 75μm I.D. Acclaim® PepMap RSLC column using gradients of 4% to 25% acetonitrile at a flow rate of 300 nL/min (solvent A: 100% H_2_O, 0.1% formic acid; solvent B: 100% acetonitrile, 0.1% formic acid) prior to MS^n^ analysis. For each MS^n^ acquisition, duty cycles consisted of one full Fourier transform scan mass spectrum (375–1500 m/z, resolution of 60,000 at m/z 400) followed by data-dependent MS^2^ and MS^3^ acquired at top speed in the Orbitrap and linear ion trap, respectively. Ions detected in MS^1^ with 4+ or greater charge were selected and subjected to CID fragmentation (NCE 23%) in MS^2^ and resulting ions were detected in the Orbitrap (resolution 30,000). The top 4 abundant ions observed in each MS^2^ spectrum with charge 2+ or greater were selected and fragmented in MS^3^ using CID (NCE 35%) and detected in the linear ion trap in ‘Rapid’ mode.

MS^3^ spectral data were extracted from .raw files into .mgf format using MSConvert (Protein Wizard 3.0.21288). Extracted MS3 spectra was subjected to Protein Prospector (v.6.3.3) for database searching using Batch-Tag against a randomly concatenated human protein database (SwissProt.2021.10.02, 20387 entries). The mass tolerances were set as ±20 ppm for parent ions and 0.6 Da for fragment ions. Trypsin was set as the enzyme with three maximum missed cleavages allowed. Cysteine carbamidomethylation was selected as fix modification. A maximum of three variable modifications were also allowed, including methionine oxidation, N-terminal acetylation, and N-terminal conversion of glutamine to pyroglutamic acid. Three defined DSSO cross-linked modification on uncleaved lysines, including alkene (C_3_H_2_O, +54 Da), thiol (C_3_H_2_SO, +86 Da) and sulfenic acid (C_3_H_4_O_2_S, +104 Da) were also selected as variable modifications. MS^n^ data were integrated via in-house software xl-Tools to identify cross-linked peptide pairs.

### LC-MS/MS analysis and protein identification/label-free quantitative analysis

Digested peptides were subjected to LC MS/MS analysis using an UltiMate 3000 UHPLC (Thermo Fisher Scientific) coupled on-line to an Orbitrap Fusion Lumos mass spectrometer (Thermo Fisher Scientific). Reverse-phase separation was performed on a 50cm x 75μm I.D. Acclaim® PepMap RSLC column. Peptides were eluted over an 87 min gradient of 4% to 25% acetonitrile at a flow rate of 300 nl/min (solvent A: 100% H2O, 0.1% formic acid; solvent B: 100% acetonitrile, 0.1% formic acid). Each cycle consisted of one full Fourier transform scan mass spectrum (375–1800 m/z, resolution of 60,000 at m/z 400) followed by data-dependent MS/MS acquired for 3 s at top speed in the linear ion trap with HCD 25% NCE. Target ions already selected for MS/MS were dynamically excluded for 60 sec after being selected twice.

MS raw data was analyzed by Maxquant software (ver 1.5.3.12) for protein identification and Perseus software for label-free quantitative analysis (46, 47). For protein identification, Cys cabamidoacetamide was specified as a fixed modification, Met oxidation, hydroxyproline and protein N-terminal acetylation was specified as variable modifications with trypsin as the proteolytic enzymes allowing for a maximum of 2 missing cleavages. The data was searched against Uniprot human reference proteome database (downloaded 02/23/2021 with 75776 sequences) concatenated with reversed sequence database as decoy. Mass tolerance of FTMS for Orbitrap was set at 20 ppm for the first search and 4.5 ppm for the main search and the mass tolerance of ion trap MS/MS was set at 0.5 Da. Proteins were identified with a false discovery rate of 1% at the levels of protein, peptide and modification site. Match-between-run was enabled. IBAQ values were calculated by Maxquant for label-free quantification analysis considering only unmodified peptides as well as peptides with protein N-terminal acetylation.

For label-free quantification analysis in Perseus, only the raw files of the triplicate PHD2 interactome analysis were included (V1 to V3 for vector control group, P1 to P3 for PHD2 group, D1 to D3 for PHD2 group with DMOG treatment). Protein groups identified only by site were first filtered out. IBAQ values of protein groups were log10 transformed. Raw data for triplicate analysis of control, PHD2 expression and PHD2 expression with DMOG treatment were grouped and only protein groups quantified in all three MS raw data in at least one sample group were kept for quantitative analysis. Missing values were replaced with normal distribution of total matrix with default settings. Two-samples Student’s T-test were performed between each pair of sample groups with permutation-based false discovery rate cutoff of 0.05 and presented in volcano plots. Selected gene names of the leading protein in each protein group were highlighted in the volcano plots.

### Bioinformatic analysis

Proteins that were identified as significant interacting proteins of PHD2 with or without DMOG treatment were overlapped and presented in Venn diagrams using R package “VennDiagram”. Overlapped proteins were subjected to Gene Ontology annotation enrichment analysis using WebGestalt with corrected false discovery rate of 0.05 (57). Protein-protein interaction network was presented using online STRING webserver with interaction score cutoff of 0.3 based on full STRING network with edges representing evidence (58, 59).

### Crosslinking-mediated structural modeling

Modeling of PHD2 and HIF1α interactions based on known structures and crosslink analysis was performed with Alphalink2 using Linux clusters from Minnesota Supercomputing Institute following instructions published on Github (https://github.com/Rappsilber-Laboratory/AlphaLink2?tab=readme-ov-file) (60). Crosslinked protein domain map was generated by xiView server (61). Structural visualization was performed with ChimeraX (version 1.8) (62).

## AUTHOR CONTRIBUTION

YC and HL designed the study and supervised the experiments. AL generated cell lines. HO and CH performed proteomics sample preparation and functional analysis with western blotting. XW assisted on generating HB-tagged stable cell lines and optimizing affinity purification. XW and CY developed crosslinking workflow and performed XL-MS and LC-MS analyses. CY, CH and YC performed data analysis. YC drafted the manuscript with inputs from all authors.

## ACKNOWLEDGEMENT

We greatly appreciate the discussion and suggestions from the members of the Chen lab and helpful advice from Drs. David Bernlohr, Do-Hyung Kim, Jeongsik Yong and Douglas Mashek. YC was supported by funding from the University of Minnesota and the National Institute of Health (GM124896 to YC). This work was also supported by R35GM145249 to L.H..

