## Supporting Information for "Crosslinking-mediated Interactome Analysis Identified PHD2-HIF1α Interaction Hotspots and the Role of PHD2 in Regulating Protein Neddylation"

\# indicates equal contributions

**Supplementary Figures**

Figure S1. MS^n^ Identification of PHD2-H1Fα Cross-linked Peptides

Figure S2. Alphafold-predicted structure of full-length PHD2.

Figure S3. Illustration of domains and regions on HIF1A (Q16665) and PHD2 (Q9GZT9).

Figure S4. Predicted structural interactions between full-length PHD2 and HIF1A.

Figure S5. Label-free quantitative analysis of PHD2 interactome with or without DMOG trapping.

Figure S6. Western blotting analysis showing that PHD2-CSN complex interaction was independent of HIF1α.

Figure S7. Western blotting analysis showing that PHD2-CSN complex interaction was not affected by hypoxia conditions.

Figure S8. Western blotting analysis for the validation of PHD2-CSN complex interaction in 293T cells.

Figure S9. A list of reported Hyp sites on CSN complex in HypDB.

Figure S10. Western blotting analysis showing that DMOG treatment reduced the poly-ubiquitination of known Cullin3 targets, PHD2 and NRF2.

**Supplementary Tables**

**Table S1.** Identification of inter- and intra-crosslinking sites of PHD2/EGLN1 and HIF1α.

**Table S2.** Identification and label-free quantification of PHD2 interacting proteins. (A) A complete list of identified proteins from interactome analysis of PHD2 interactome including four replicate analysis of each group. (B) PHD2 interacting proteins that were significantly upregulated with streptavidin pulldown from HeLa cells stably expressing HTBH-tagged PHD2 vs HeLa cells expressing HTBH-vector control. (C) PHD2 interacting proteins that were significantly upregulated with streptavidin pulldown from PHD2-expressing HeLa cells with DMOG trapping vs HeLa cells expressing HTBH-vector control.

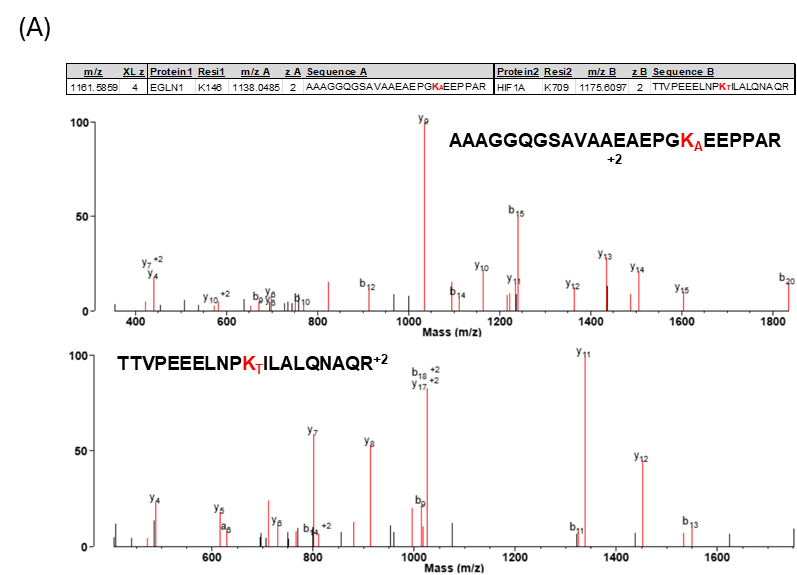

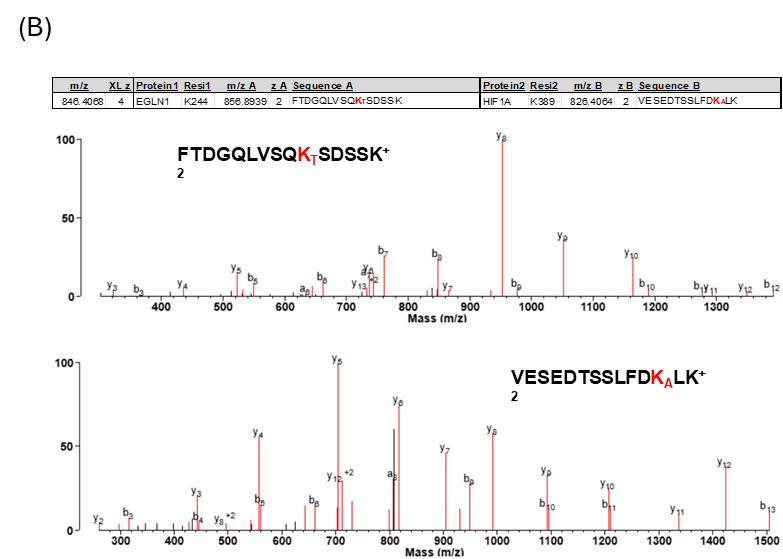

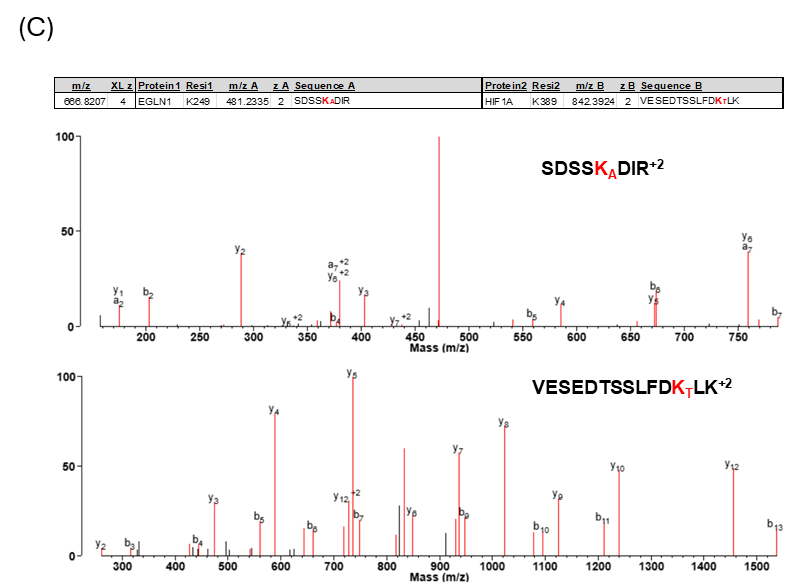

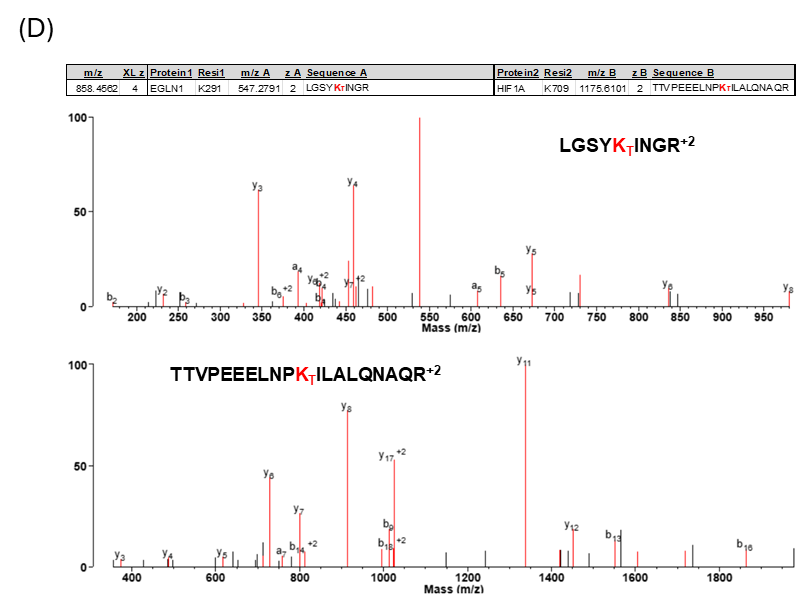

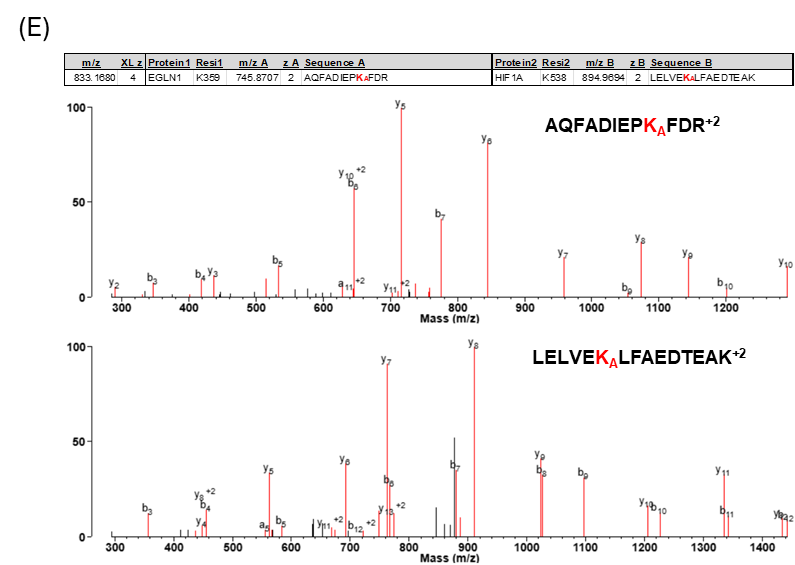

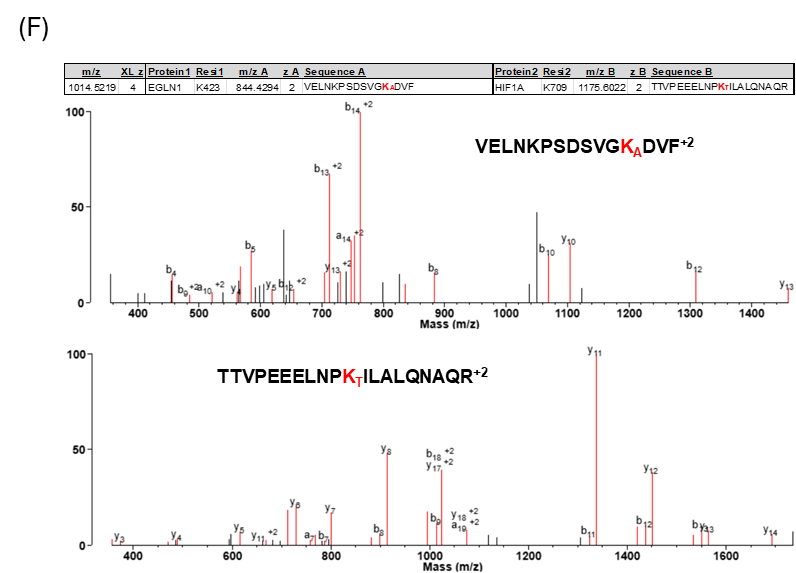

**Figure S1.** MS^n^ Identification of PHD2-H1Fα Cross-linked Peptides. (A) PHD2 K146-HIF1α K709, (B) PHD2 K244 – HIF1α K389, (C) PHD2 K249 – HIF1α K389, (D) PHD2 K291 – HIF1α K709, (E) PHD2 K359 – HIF1α K538, (F) PHD2 K423 – HIF1α K709.

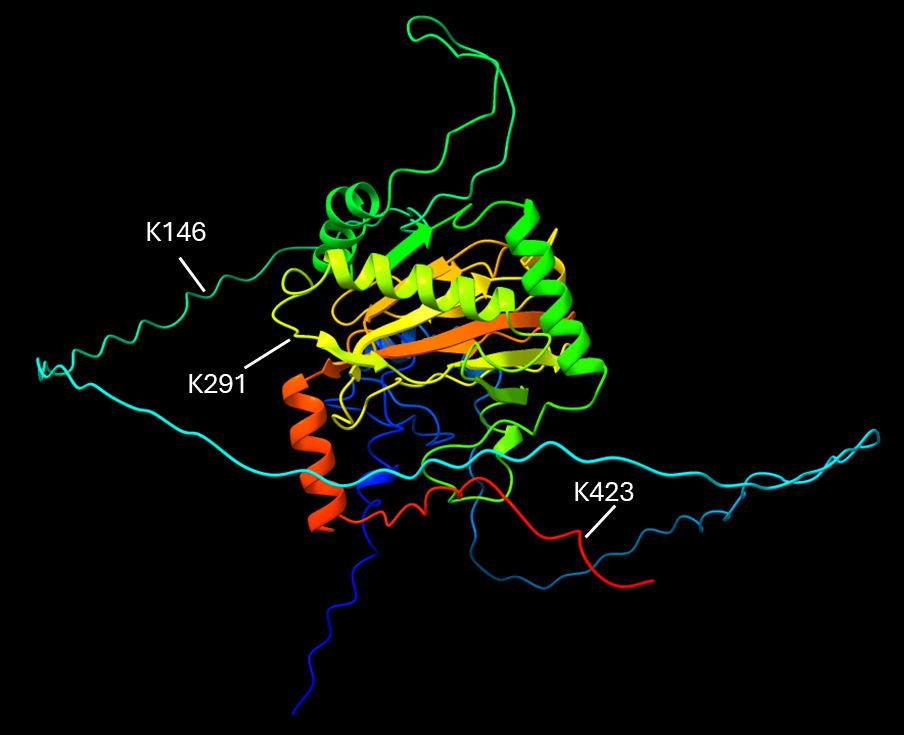

**Figure S2.** Alphafold-predicted structure of full-length PHD2. Three positions highlighted were K146, K291 and K423 on the PHD2 that were identified to crosslink with HIF1α K709.

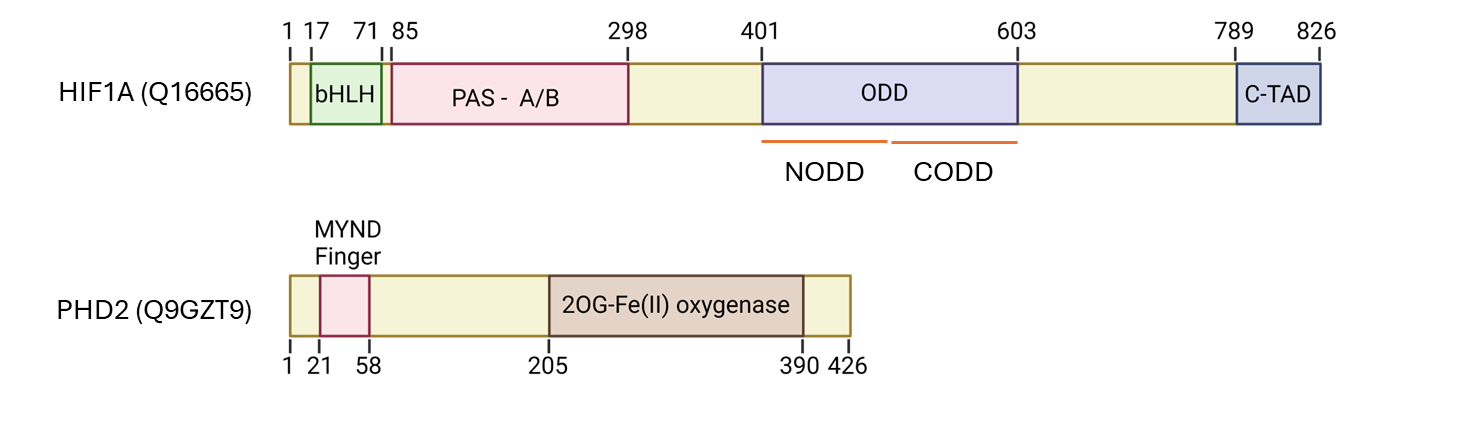

**Figure S3.** Illustration of domains and regions on HIF1α (Q16665) and PHD2 (Q9GZT9).

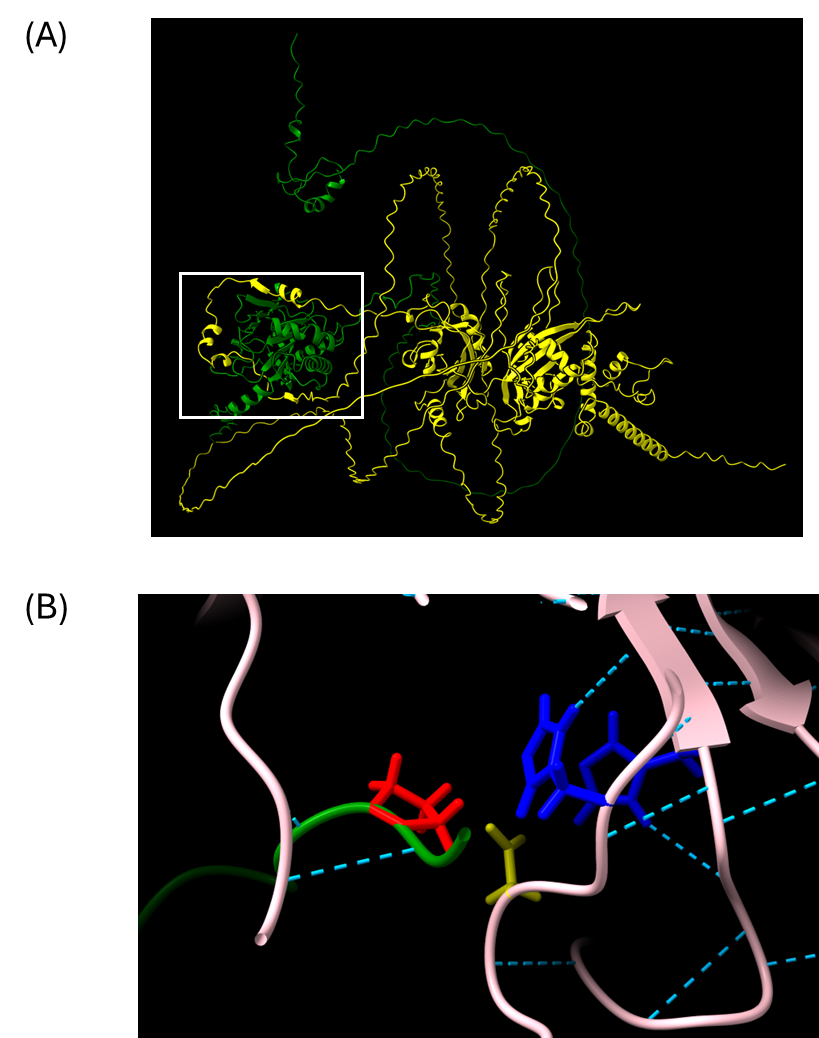

**Figure S4.** Predicted structural interactions between full-length PHD2 and HIF1α. (A) Best scored model of full-length PHD2 (in green) and HIF1α (in yellow). (B) The close positioning of HIF1α P402 (in red) with PHD2 catalytic triad H313 (in blue), D315 (in yellow) and H374 (in blue).

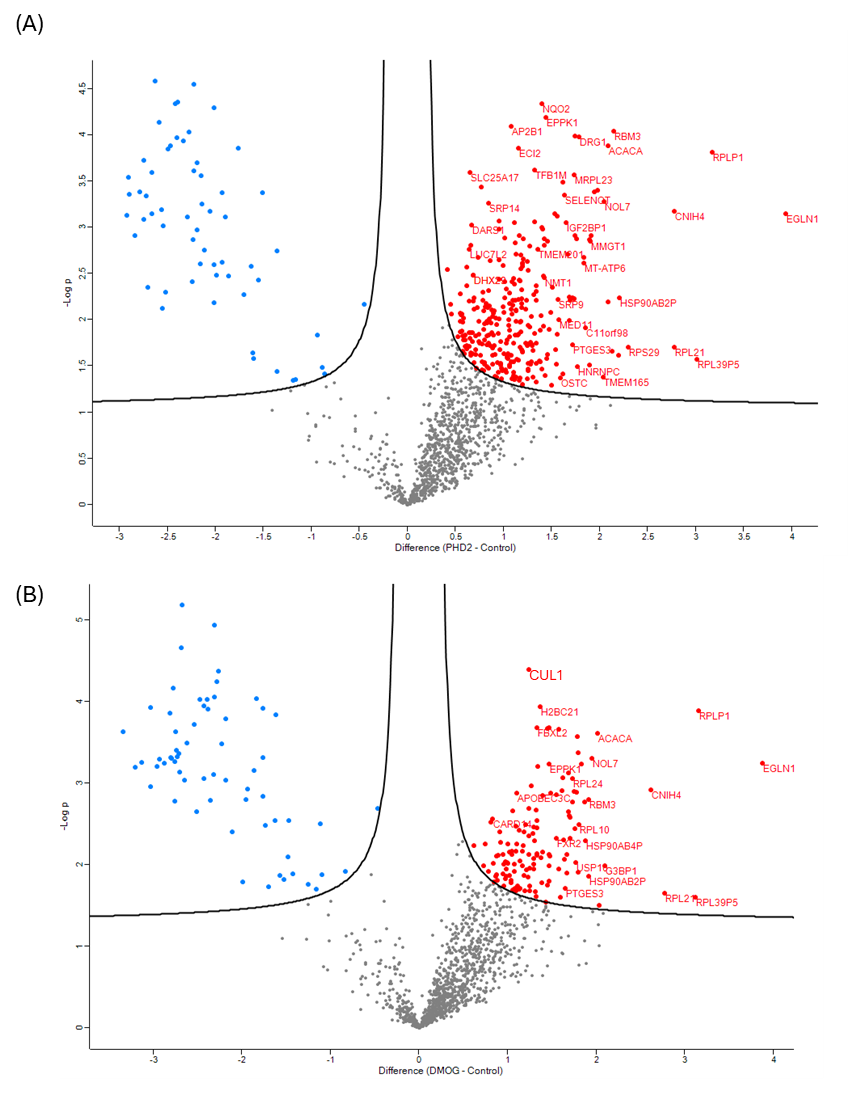

**Figure S5.** Label-free quantitative analysis of PHD2 interactome with or without DMOG trapping. (A) Volcano plot analysis of PHD2 interacting proteins comparing protein quantifications in HeLa cells expressing HTBH-PHD2 with control HeLa cells (FDR<0.05). (B) Volcano plot analysis of PHD2 interacting proteins comparing protein quantification in HeLa cells expressing HTBH-PHD2 upon DMOG treatment with control HeLa cells (FDR<0.05).

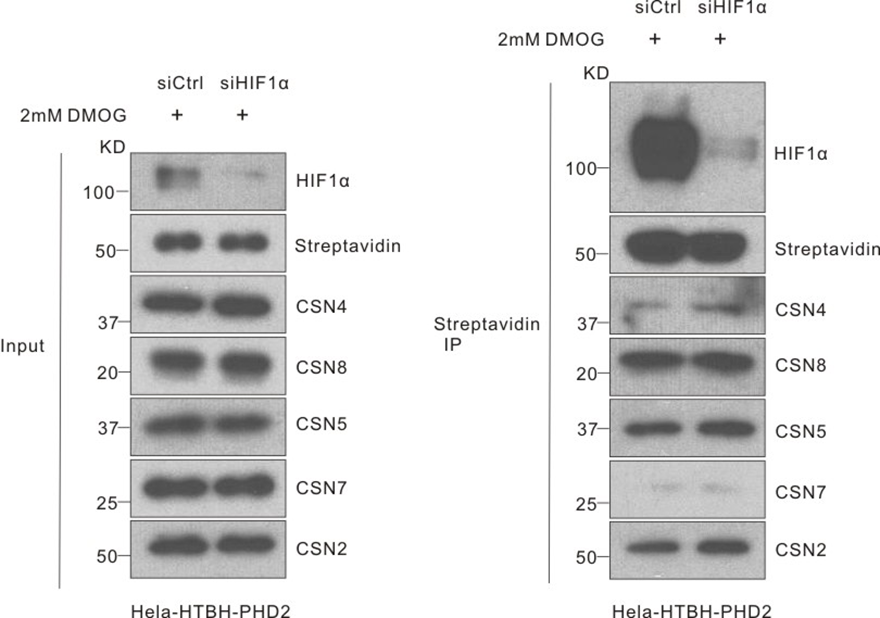

**Figure S6.** Western blotting analysis showing that PHD2-CSN complex interaction was independent of HIF1α. HIF1α was knocked down in HeLa cells that stably express HTBH-PHD2. Then, cells were treated with 2mM DMOG for 4 hours prior to lysis and streptavidin enrichment. The same amount of input and IP samples of each group were loaded for WB analysis and blotted with antibodies as indicated. The expression of HIF1α was examined as an indicator of the knockdown efficiency.

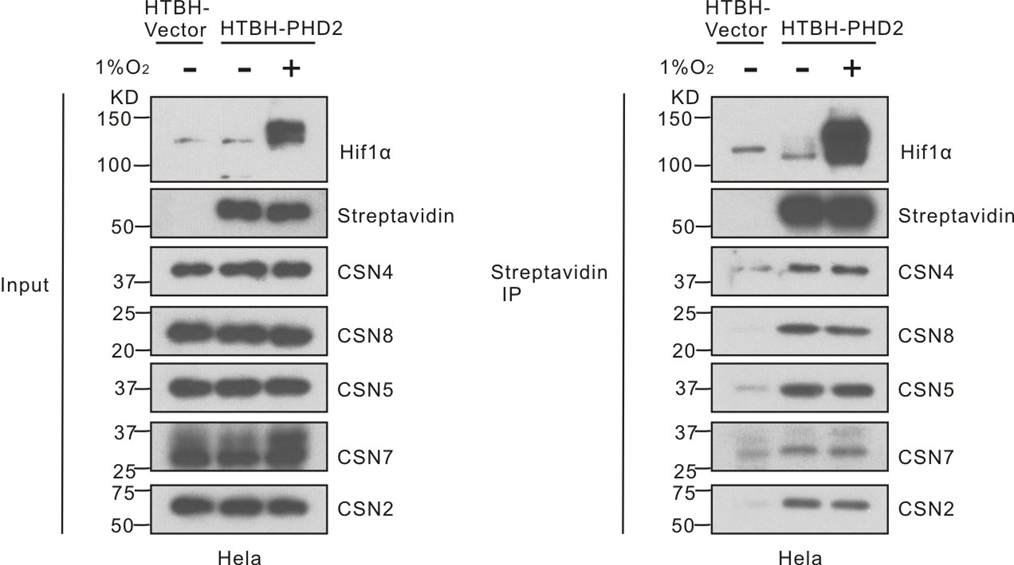

**Figure S7.** Western blotting analysis showing that PHD2-CSN complex interaction was not affected by hypoxia conditions. HeLa cells that stably express HTBH-empty vector or HTBH-PHD2 were seeded and cultured under normoxia or hypoxia for 5 hours prior to lysis and streptavidin enrichment. The same amount of input and IP samples of each group were loaded for WB analysis and blotted with antibodies as indicated. The expression of HIF1α was examined as an indicator of the hypoxia treatment.

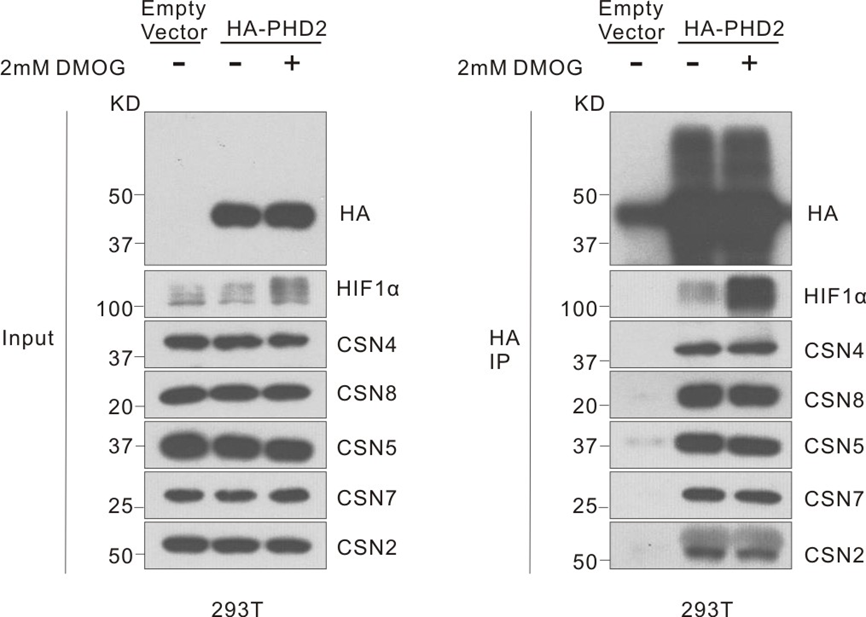

**Figure S8.** Western blotting analysis for the validation of PHD2-CSN complex interaction in 293T cells. 293T cells were transiently transfected with HA-PHD2 plasmid or empty vector for 18 hours and treated with 2mM DMOG or DMSO for the next 4 hours prior to lysis and immunoprecipitation with anti-HA antibody conjugated beads. The same amount of input and IP samples of each group were loaded for WB analysis and blotted with antibodies as indicated. The expression of HIF1α was examined as the indicator of DMOG treatment.

(A)

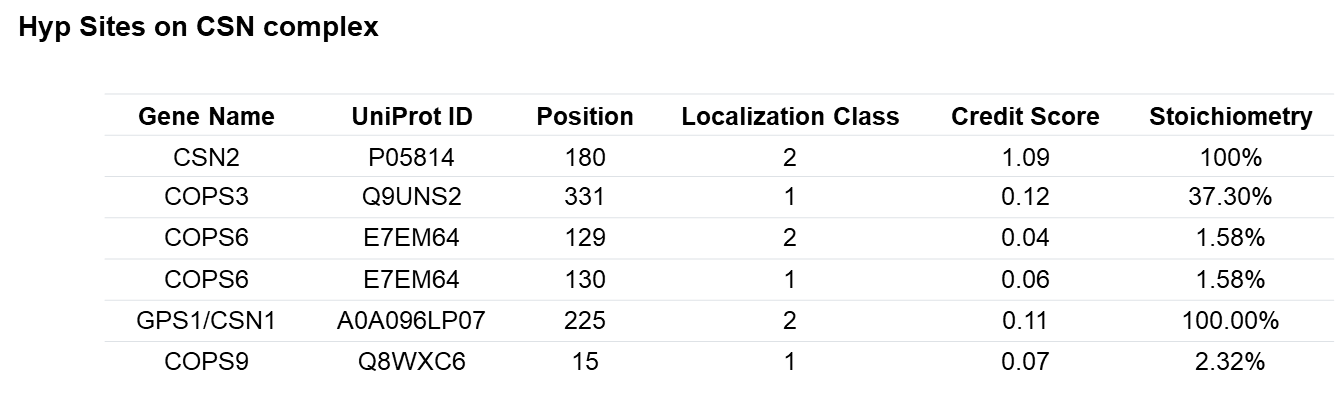

(B)

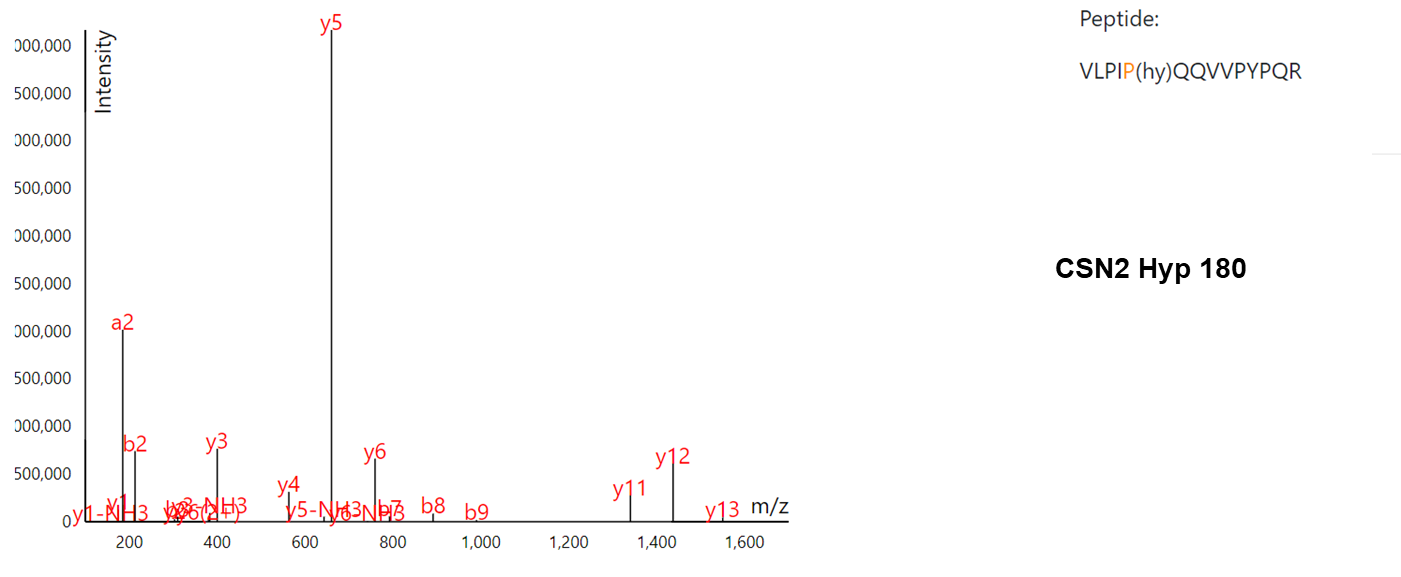

(C)

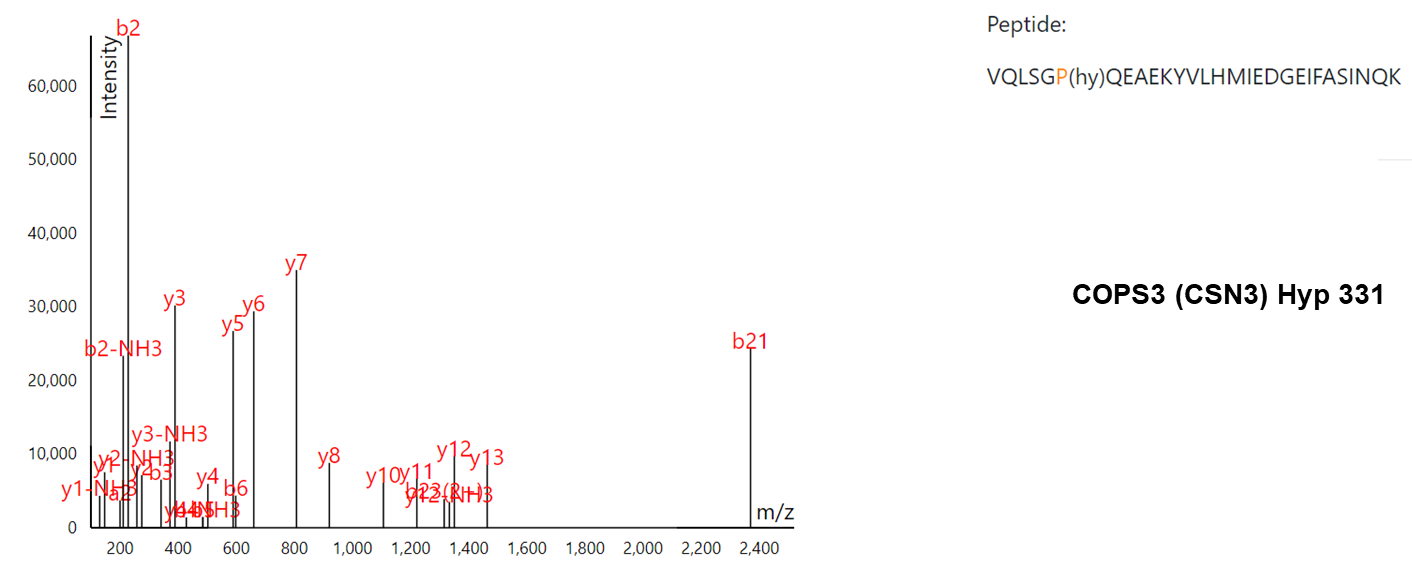

(D)

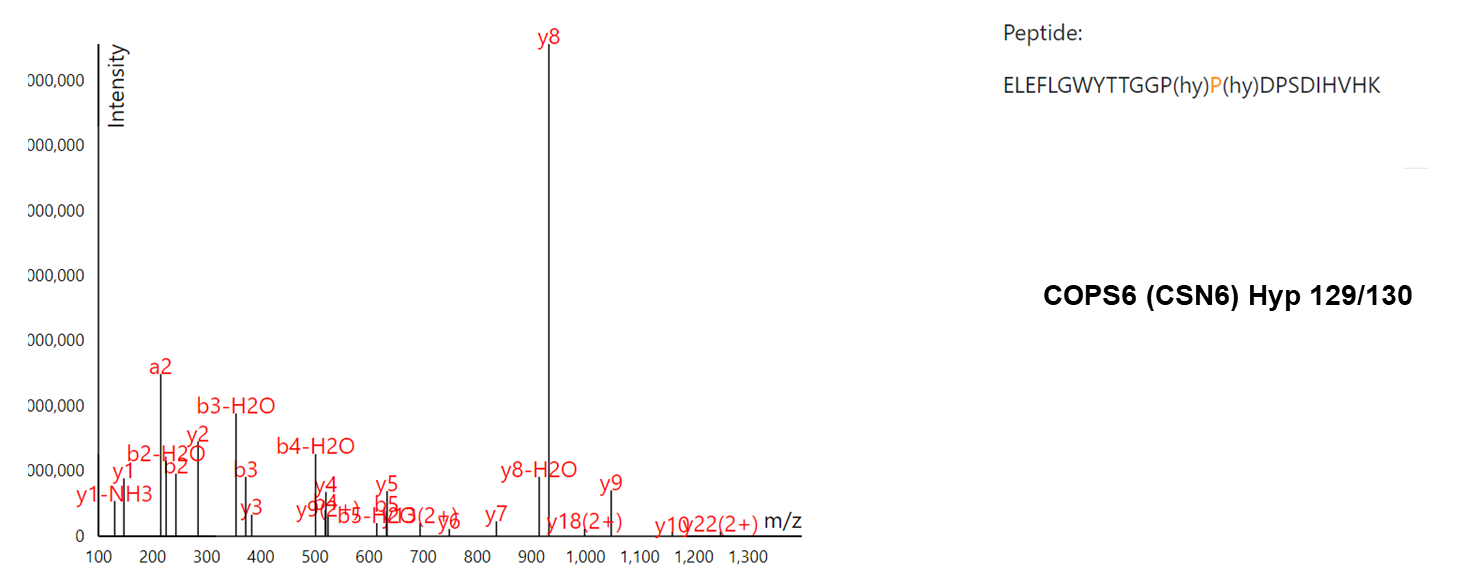

(E)

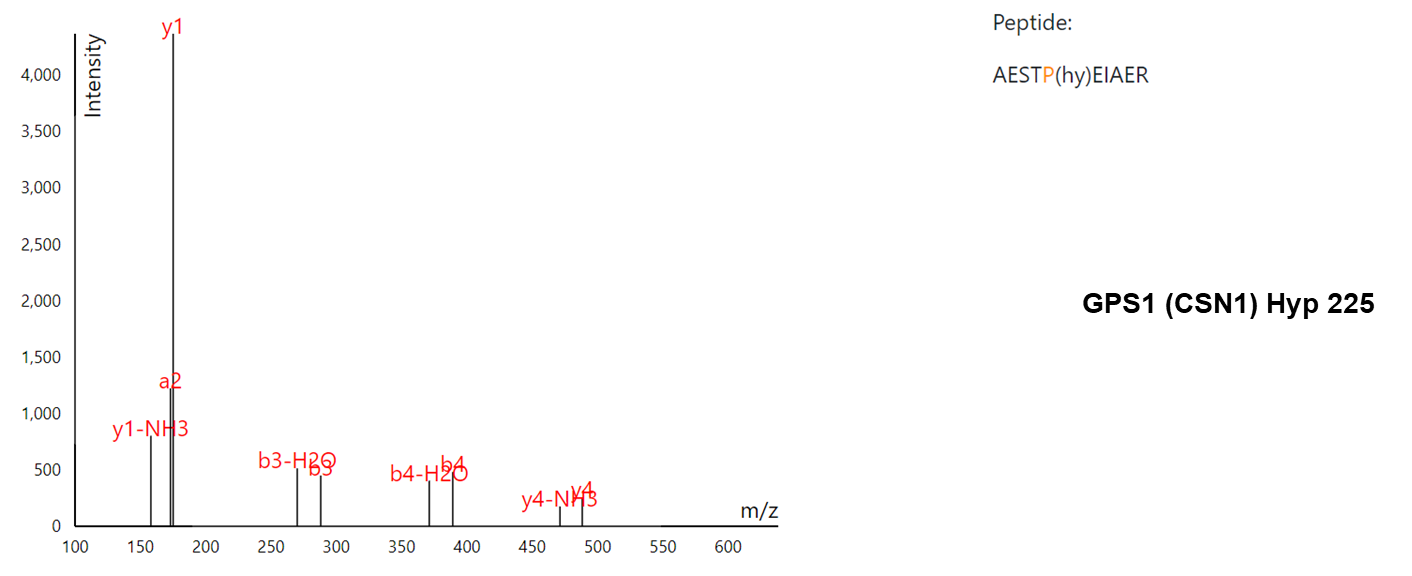

(F)

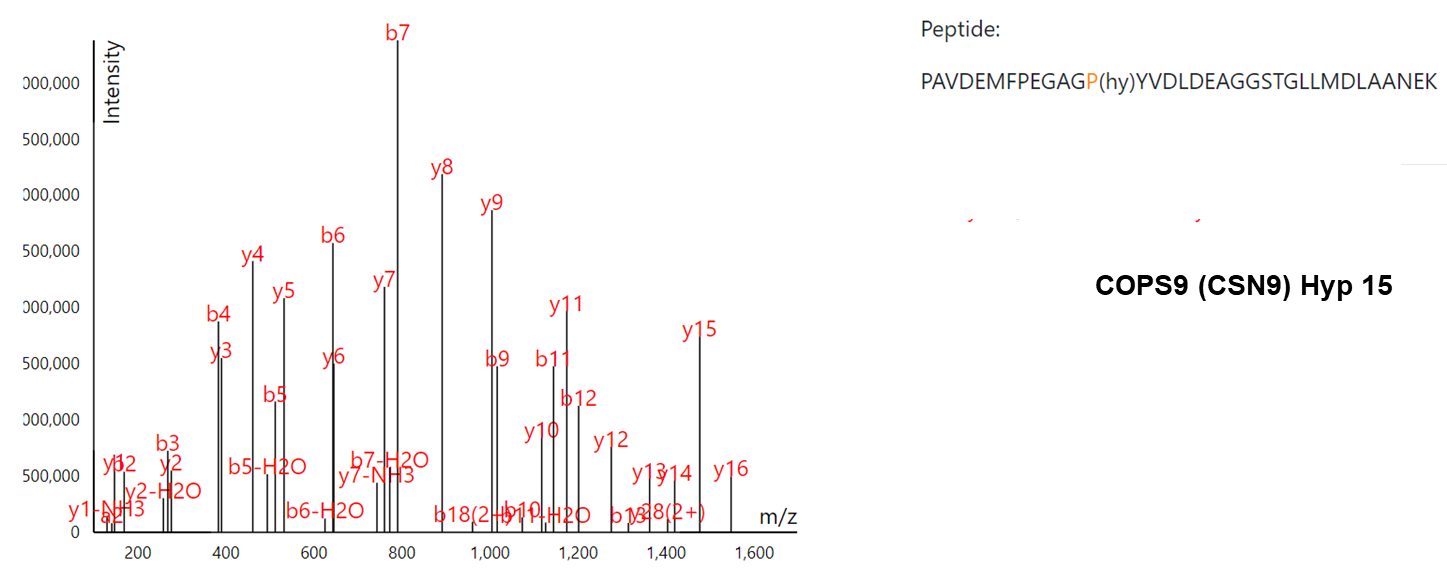

**Figure S9.** A list of reported Hyp sites on CSN complex in HypDB. (A) a table presentation of six Hyp sites reported on the members of the CSN complex in the HypDB database. Annotated mass spectra for identified Hyp peptides (B) CSN2 Hyp180, (C) COPS3 Hyp331, (D) COPS6 Hyp129/130, (E) GPS1/CSN1 Hyp225, (F) COPS9 Hyp15. “(hy)” indicated hydroxyproline sites. a, b, y ions represented different types of peptide backbone fragmentations.

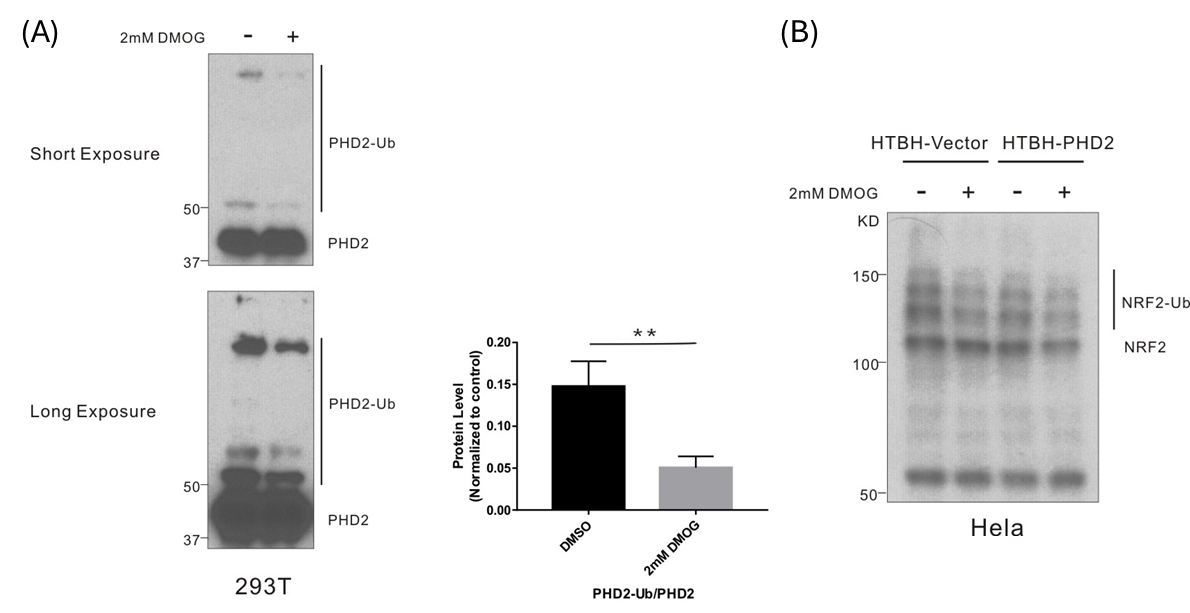

**Figure S10.** Western blotting analysis showing that DMOG treatment reduced the poly-ubiquitination of known Cullin3 targets, PHD2 and NRF2. (A) 293T cells were seeded and treated with DMSO or 2mM DMOG for the next 24 hours and cultured under normoxia condition. Cells were lysed for WB analysis with anti-PHD2 antibody. n=4, *P<0.05, ** P<0.01, ***P<0.001. (B) Hela cells stably expressing HTBH-empty vector or HTBH-PHD2 were seeded and treated with DMSO or 2mM DMOG for 24 hours under normoxia, then cells were harvested and lysis for WB analysis and blotted with anti-NRF2 antibody.
